# The Cone Method: Inferring Decision Times from Single-Trial 3D Movement Trajectories in Choice Behavior

**DOI:** 10.1101/2020.08.01.232314

**Authors:** Philipp Ulbrich, Alexander Gail

## Abstract

Ongoing goal-directed movements can be rapidly adjusted following new environmental information, e.g. when chasing pray or foraging. This makes movement trajectories in go-before-you-know decision-making a suitable behavioral readout of the ongoing decision process. Yet, existing methods of movement analysis are often based on statistically comparing two groups of trial-averaged trajectories and are not easily applied to three-dimensional data, preventing them from being applicable to natural free behavior. We developed and tested the *cone method* to estimate the point of overt commitment (POC) along a single two- or three-dimensional trajectory, i.e. the position where movement is adjusted towards a newly selected spatial target. In Experiment 1, we established a “ground truth” data set in which the cone method successfully identified the experimentally constrained POCs across a wide range of all but the shallowest adjustment angles. In Experiment 2, we demonstrate the power of the method in a typical decision-making task with expected decision time differences known from previous findings. The POCs identified by cone method matched these expected effects. In both experiments, we compared the cone method’s single trial performance with a trial-averaging method and obtained comparable results. We discuss the advantages of the single-trajectory cone method over trial-averaging methods and possible applications beyond the examples presented in this study. The cone method provides a distinct addition to existing tools used to study decisions during ongoing movement behavior, which we consider particularly promising towards studies of non-repetitive free behavior.

## Introduction

While interacting with our environment, we often need to make fast choices about our upcoming path of movement while already being engaged in an ongoing action. When we hurry through the train station to catch our connection, we have to dodge other passengers, the same way soccer players need to quickly decide which direction to dribble or pass the ball in response to the movement of their opponents, or hunting animals have to pick their prey from a fleeing herd. Being able to make such online choices is the result of a continuous sensorimotor integration process, as we constantly take in new information to dynamically select and adapt our actions (see Pezzulo & Cisek, 2016 for review). Consequently, decision-making and the preparation and control of action are considered to be parallel and interlinked rather than separate and serial processes, both, at the behavioral (e.g. Morel et al., 2017) and neural level (e.g. Cisek et al., 2005; Klaes et al., 2011; Pastor-Bernier & Cisek, 2011; Suriya-Arunroj & Gail, 2019; see also Gallivan et al., 2018; Wispinski et al., 2020 for review). Decision-related online movement adjustments have been demonstrated in movements ranging from timescales of more than a second (e.g. Cheng & González-Vallejo, 2017), over rapid reaches lasting only a few hundred milliseconds (e.g. Chapman et al., 2015), down to the fastest correctional responses to sudden perturbations (Carroll et al., 2019; Nashed et al., 2012, 2014). Here we want to introduce a method that allows us to estimate the timing of decision-related trajectory adjustments towards a spatial target from an individual movement trajectory.

One class of behavioral paradigms that capitalizes on our ability to online select spatial targets are commonly referred to as *go-before-you-know* (Gallivan & Chapman, 2014) or mouse tracking tasks (Freeman et al., 2011; Spivey et al., 2005). In both cases, tight time constraints require subjects to initiate their movements prior to knowing which of multiple potential targets to select. Thus, subjects need to adjust their movement mid-flight and the resulting trajectories serve as online behavioral readout of cognitive processes such as sensory processing and decision-making as they unfold during the movement (Chapman et al., 2014; Dotan et al., 2019; Scherbaum et al., 2010). Conversely, conventional “first decide, then act” reaction time paradigms provide all necessary information prior to the behavioral response and the latter (e.g. a button press) only serves as confirmation of the already finished target selection process without the opportunity for online revisions of the choice. However, reaction time paradigms by design have the advantage that they provide an inherent time measure for the termination of a decision process (the reaction time), which is not easily obtained in online decisions during ongoing movements.

Our newly developed *cone method* aims at combining the advantages of online decision-making paradigms with the ability of estimating decision times of individual choices directly from movement trajectories. The method continuously compares the current movement direction with the range of possible directions aiming at the finite-size target. We assess how reliably the cone method allows to identify straight-to-target movement, i.e. cases in which a decision is made prior to movement start and, in the other cases, to estimate when a decision occurred along a movement trajectory. While the cone method was conceived with behavioral paradigms in mind that possess a segmented trial-by-trial structure, its independence from a defined movement onset point makes it applicable to more naturalistic paradigms that utilize continuous movements without a trial structure, such as chasing pray or foraging.

## Methods

### Conceptual Experimental Design

We conducted two experiments. The first experiment aimed at establishing “ground truth” data for tuning the cone method. The second experiment created a test scenario under the realistic conditions of a choice experiment with predictable outcome. In the first experiment, subjects performed go-before-you-know reaches towards instructed targets. During the initial segment of the movement along a predefined trajectory (*movement corridor*), one of eight possible locations was visually indicated as the target. Subjects could only curve the movement towards the laterally offset target within a predefined region in space (*via-sphere*) along the movement corridor. We thereby controlled when the online adjustment of movement had to happen (*point of commitment* [POC] to the target) and used this information as ground truth for confirming how well our newly developed cone method was able to recover these POCs within their known spatial constraints. We hypothesized that the cone method’s performance would depend on the steepness of the angle at which the movement had to be adjusted towards the target (*adjustment angle*), with larger adjustment angles leading to better performance due to the more easily detectable curvature of the movement. Therefore, we varied the distance of the via-sphere from the hand starting position and the lateral offset of the target, which required the subjects to produce a wide range of adjustment angles.

In Experiment 2, we used a similar layout as in Experiment 1, but asked subjects to choose between two continuously visible targets of different value in order to maximize their reward. Subject were free in how to approach their chosen target, i.e. no via point constrained the movement, but they had to be fast. In each trial, subjects either decided between gaining tokens and a neutral (+/-0 token) outcome (*gain trial*) or losing tokens and a neutral outcome (*loss trial*). Previous results showed that when cueing the target-associated value information at different *stimulus onset asynchronies* (SOA) relative to the go cue, the value cue in the loss trials had to be presented on average 100ms earlier compared to the gain trials to elicit similar trajectories (Chapman et al. 2015). The idea of Experiment 2 is to use this strong effect on choice latency to compare our cone method with an established trial-averaging trajectory measure in an actual choice experiment.

### Setup

Both experiments were conducted in a 3D augmented reality (3D-AR, Figure 1A) haptic reach setup (Morel et al., 2017). Subjects performed reaching movements using a parallel-type haptic manipulator (Delta.3, Force Dimension, Nyon, Switzerland). The manipulator was connected to a computer running custom software (C++, OpenGL), responsible for task control, incl. visual stimulus generation, hand position recording (manipulator handle position sampled at 2 kHz), and task event recording (digital IO).

**Figure 1.**
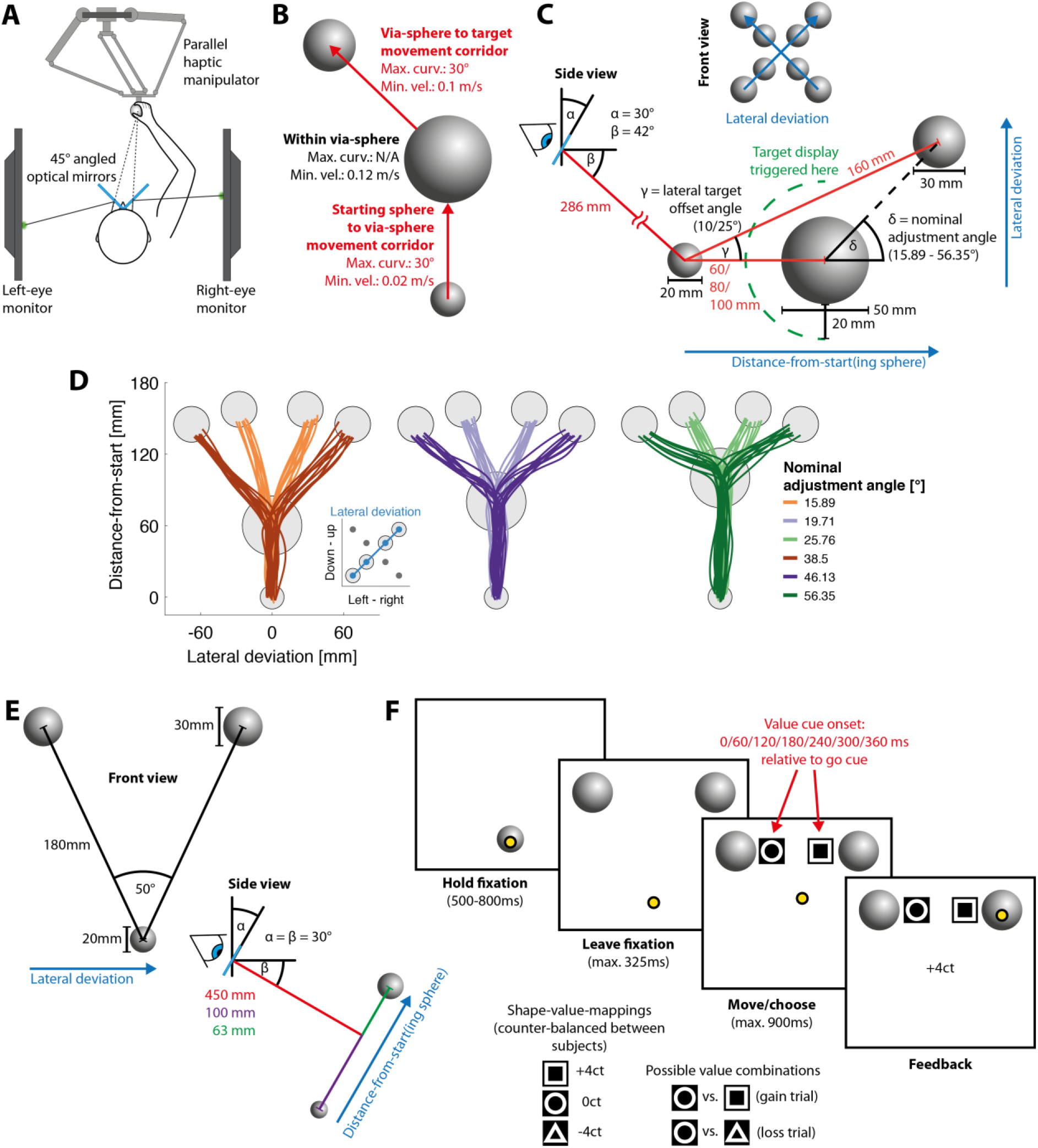
3D Augmented reality setup and behavioral paradigms. **A:** Augmented reality setup. Subjects performed reaching movements by grasping and moving a haptic manipulator. Visual stimuli such as cursor and reaching targets were viewed through a mirror stereoscope and thereby projected directly into the 3D manipulator’s workspace. **B:** Experiment 1, behavioral paradigm. Subjects moved the handle from the starting sphere (bottom) through the via-sphere (middle) to the target sphere (top) while adhering to the maximum curvature and minimum velocity constraints. **C:** Experiment 1, stimulus arrangement. For ergonomic reasons the manual workspace was below eye level. For this, the optical mirrors were tilted down by 30° (α) such that subjects viewed the stimuli diagonally from the top (β) and performed the movements away from the body. The target was displayed only once the cursor was close enough (green dashed line) to the via-sphere to prevent anticipatory curving of the movement prior to entering the via-sphere. For 2D illustration purposes, the trajectories were projected onto the lateral deviation axis while retaining the original Z-axis (“distance—from-start”). **D:** Experiment 1, single subject example trajectories for target directions 45° and 225°. **E:** Experiment 2, stimulus arrangement (bottom: starting sphere; top: potential reach targets). The mirror tilt, identical to Experiment 1, defined the orthogonal workspace plane on which the starting- and target spheres were placed. **F:** Experiment 2, example trial, possible mappings of value cue symbols to trial outcomes, and possible value cue combinations. Subjects reached from the starting sphere to one of the two targets and were asked to select the target based on the value cue in order to maximize the outcome of the trial. The value cue contrast in this figure is inverted compared to its actual appearance.

The 3D-AR environment consisted of two computer monitors (BenQ XL2720T, 590mm × 338mm screen size, 60 Hz refresh rate, 47mm viewing distance, driven with a Matrox DualHead2Go Display Port Splitter) that were viewed through a pair of semitransparent mirrors, tilted 45° relative to the screens. Subjects only viewed one screen per eye, which allowed for the creation of stereoscopic 3D images perceived as directly projected into the manipulator’s workspace. This means that all movement-related stimuli such as movement starting points and targets were directly presented at their supposed physical location. The position of the manipulator’s handle was represented in the AR as yellow sphere cursor (d = 6mm) at its actual physical location. Display and manipulator latencies were fully compensated by a forward prediction using a Kalman filter with position, speed, and acceleration as state variables to synchronize the movement of the handle and the cursor. The haptic manipulator was mounted approx. at chest height to allow for comfortable operation. Consequently, the monitors and the mirror were additionally tilted by 30° to lower the 3D representation into the manipulator’s workspace. (Figure 1C, angle α).

The 3D-AR environment was calibrated by making the physical handle position coincide with multiple sequentially presented visual targets. This manually adjusted handle position was used to compute a manipulator-to-display transformation matrix. This calibration was further adjusted for each subject by setting the location and projection matrix of the virtual openGL cameras according to the subject’s interpupillary distance.

### Participants

In Experiment 1, eight lab-internal subjects (age *M*±*SD* = 26.4±4, 3 female, 1 left-handed, all normal or corrected-to-normal vision), including the first author of this study, participated without receiving monetary compensation. Due to the straightforward nature of the experiment, subject naivety was not required, and subjects were instructed verbally by the experimenter.

In Experiment 2, six subjects (age *M*±*SD* = 23.3±3.4, 3 female, all right-handed, all normal or corrected-to-normal vision) participated, recruited via the university’s notice board. All subjects were naive to the purpose of the task and had not participated in similar experiments before. While the behavioral paradigm of Experiment 2 matches elements of Chapman et al. (2015), the purpose of this study is not to replicate the previous study now with a 3D manipulator task, but rather to exploit a known effect of choice latency on ongoing movement trajectories to demonstrate the cone method’s capability to capture ongoing decisions. For this purpose, six subjects were enough to express the effect of a gain- versus loss framing on the trajectories and at the same time to provide sufficient variability in movement patterns. The subjects received a base remuneration of 8€ per hour, with the experiment lasting on average 1.5-2 hours. Based on their choices and performance during the experiment, subjects could either gain an overall bonus or suffer an overall loss. Bonuses were added to the base remuneration in full. Losses, where applicable, would have been capped such that the net remuneration does not fall below 6€ per hour. Subjects were unaware of the latter and there was no need to apply this rule. On average, subjects received a bonus of 2.53€ (*SD* = ±0.87).

In both experiments, subjects gave their written informed consent prior to participation. Experiments were in accordance with institutional guidelines for experiments with humans, adhered to the principles of the Declaration of Helsinki, and were approved by the ethics committee of the Georg-Elias-Mueller-Institute for Psychology at the University of Goettingen.

### Experiment 1: Simulated Online Decisions via Constrained Movement Trajectories

In Experiment 1 (Figure 1 B & C), subjects were asked to move the handle with their dominant arm from a starting sphere (d = 20mm) through a via-sphere (d = 50mm) to a target sphere (d = 30mm) in 3D space. Subjects were required to maintain a certain minimum velocity during each stage of the movement (starting sphere to via-sphere: 0.02m/s; within via-sphere: 0.12m/s; via-sphere to target: 0.1m/s). Outside the via-sphere, the angular deviation of movement was constrained to maximally 30°; i.e. the difference between the momentary movement direction and the initial movement direction when leaving the starting sphere and via-sphere, respectively, was not allowed to exceed this threshold (movement corridor). These constraints required subjects to perform fluent, non-intermitted movement adjustments towards the target within the experimentally defined via-sphere. The display of the target was triggered when the cursor fell below a 20mm distance from the edge of the via-sphere (median visibility onset = 43ms later; 5^th^ and 95^th^ percentiles = 36 and 51ms, respectively). The median target onset time relative to entering the via-sphere additionally depended on the cursor speed between surpassing the 20mm threshold and entering the via-sphere, and was then −59ms (5^th^ & 95^th^ percentile: −29ms & −100ms). This timing allowed subjects to correct their movement towards the target within the via-sphere while at the same time adhering to the minimum velocity constraints. We did not display the target earlier to avoid anticipatory movement curvature before entering the via-sphere to the amount that the 30° limit of the movement corridor would allow. Since the minimum delay to which motor corrections in response to visual stimuli can occur is around 110ms (e.g. Brenner & Smeets, 1997; Carroll et al., 2019), our subjects were not able to respond to the early onset of the target before entering the via-sphere. Unreported pilot data showed that a later onset of the target, constraining angular deviations outside the via-sphere to less than 30°, or choosing a smaller via-sphere diameter made the task too difficult.

Subjects viewed the starting sphere from diagonally above, at an elevation of 42° below the horizontal viewing direction (Figure 1C, angle β), corresponding to 12° below the straight ahead viewing direction of the 30°-tilted setup, and at 286mm distance (both measured relative to the optical mirrors; Figure 1C). To enforce different POCs, as well as differently sized angular adjustments of the movement trajectories when starting to aim at the target, the via-sphere was located 60, 80, or 100mm away from the starting sphere (center-to-center) on the Z-axis, i.e. perpendicular to the subjects’ fronto-parallel plane. The distance from the starting sphere to the target sphere was 160mm. To sample the range of differently sized angular adjustments more densely, the target sphere was additionally laterally offset either 10° or 25 ° relative to the Z-axis (Figure 1C, angle γ). This resulted in a nominal via-sphere-to-target adjustment angle (Figure 1C, angle δ) between 15.89 and 56.35°, depending on starting-sphere to via-sphere distance and the lateral offset of the target (Figure 1D). The target sphere was located at one of four oblique directions within one of two planes (depending on the lateral offset of the targets) fronto-parallel to the subjects, spaced 90° apart, starting at 45° from the horizontal (Figure 1C, top). For illustration purposes and certain analyses, we projected the 3D movement trajectories onto a plane, defined by the Z-axis (Figure 1C *distance-from-start*) and one of two vectors orthogonal to the Z-axis, depending on the target location (Figure 1C, *lateral deviation*).

The experiment was comprised of 3 (via-sphere distance) × 2 (target sphere lateral offset angle) × 4 (target direction) = 24 conditions, which had to be successfully completed 10 times each. Conditions were drawn randomly such that, at each time during the experiment, the difference in number of successfully completed trials between each condition did not exceed one. Failed trials were not automatically repeated immediately but placed back into the pool of conditions from which the randomization algorithm drew. Additionally, subjects received verbal onscreen feedback that informed them about the type of error they made (“too slow”, or “too curved”).

Prior to the actual experiment, subjects were successively trained on two easier versions of the task (120 successful trials each). In both training sets, the final reach target was already displayed at movement onset. Additionally, in the first training set, the maximum curvature requirements outside the via-sphere were relaxed to 45° and the minimum velocity requirements were lowered to 0.02m/s for the entire movement.

### Experiment 2: Go-Before-You-Know Decision-Making Between Monetary Gains and Losses

In Experiment 2 (Figure 1 E & F), subjects performed go-before-you-know reaches towards two potential targets. Each target was associated with a different monetary outcome and subjects freely chose between the two on each trial. Subjects performed their movements from a starting sphere (d = 20mm) towards one of two targets (d = 30mm), located 180mm (center-to-center) away from the starting sphere and spaced 50° apart (Figure 1E). The location of the starting sphere and both targets was kept constant throughout the experiment. The lateral deviation – distance-from-start plane that was defined by these three stimuli was tilted 30° relative to the subjects’ fronto-parallel plane (Figure 1E, angle α). This practically resulted in a 2D task, unlike Experiment 1. Subjects were nonetheless free to guide their movements, as we did not physically constrain the movements of the manipulator handle to the task plane. Again, we projected the resulting trajectories on a plane spanned by the lateral deviation and the distance-from-start vectors. Subjects started each trial (Figure 1F) by moving the cursor into the starting sphere. After 500-800ms an auditory go cue (440 Hz, 100ms) indicated that the movement had to be initiated within 325ms. The onset of the value cue (black triangle, circle, or square on a square white background, edge length = 20mm, appearing next to the target) was triggered at an SOA of 0 - 360ms in steps of 60ms relative to the go-cue Consequently, we added 43ms to each SOA in all analyses and figures. This value corresponds to the median latency target visibility measured in Experiment 1. Subjects were supposed to pick the economically best target based on the value cue mid-flight and finish the movement within 900ms. Upon successfully acquiring one of the two targets, the amount of money gained/lost was provided as written onscreen feedback. Trials were aborted when subjects left the starting sphere too early or too late, slowed down during the movement below 0.0275m/s, or did not acquire one of the targets within 900ms after movement onset.

The three different value cue symbols were assigned a value of +4 € cent, −4 € cent or 0 € cent, respectively, which was constant throughout the experiment. The value cue-outcome mapping was counterbalanced across subjects. In each trial, one target was always associated with neutral outcome, while the other target was either associated with a gain or a loss (Figure 1F). Consequently, subjects were either supposed to decide *for* the gain target, or *against* the loss target. The aggregated net outcome was added to the subject’s base remuneration.

### Data Analysis

#### Data preparation and conventions

All data analyses and visualization were carried out with Matlab 2015b and the gramm plotting toolbox for Matlab (Morel, 2018). We obtained the movement velocity by differentiating the raw position data. To remove high frequency noise, we filtered both, the position and velocity data (zero-phase filter [Matlab function **`filtfilt`**] using a 4^th^ order Butterworth low pass filter with a 12 Hz cut-off frequency [Matlab function **`butter`**]). We resampled the filtered position and velocity data at 200 Hz using cubic spline interpolation (Matlab function **`interp1`**). For all analyses, movement start was defined as the first data point outside the starting sphere and the end of the movement as the last data point outside the target sphere. All trajectories were truncated accordingly.

#### Cone method rationale and application

We developed the cone method (Figure 2) as an algorithm to estimate from which point onwards a decision to move to a specific spatial target is visible in a single movement trajectory (point of overt commitment as recovered by the cone method, POC^cone^). For each location along a trajectory, along the distance-from-start vector (Figure 1), we define a cone with its tip at the current location and its base being defined by the maximum-diameter intersection of the spherical target area. This means, the curved surface area of the cone is defined by the tangential directions from cone tip to target circumference (see Figure 2A for a 2D representation). In effect, for each location on the trajectory this cone describes the entirety of directions aimed anywhere at the target. As long as the current direction stays outside the cone, the movement is not directed towards the target. Yet, the decision to move towards the target might have become overt through a corresponding curvature of the trajectory before entering the cone. Our cone methods accounts for this possibility and, accordingly, the POC^cone^ is defined as follows: A decision to move towards the target becomes overt in the trajectory at the earliest when the current movement direction for the first time starts to be adjusted towards the cone (Figure 2, purple marker), i.e. when the angle between the current direction and the closest direction along the cone’s surface (Figure 2B) is decreasing. This decrease has to be monotonic until the current direction stays inside the cone (Figure 2, brown marker).

**Figure 2.**
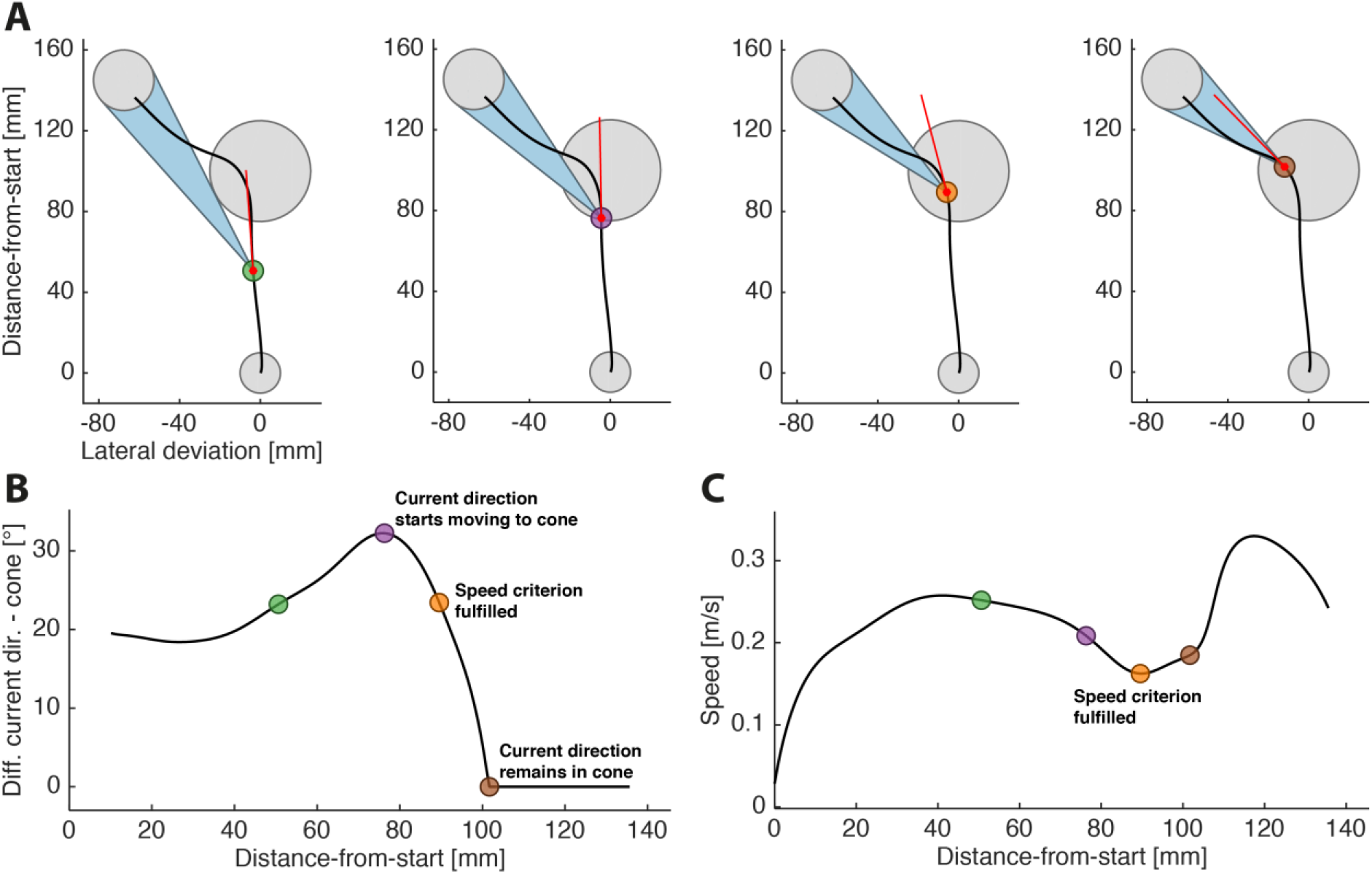
Cone method rationale. **A:** 2D projection of an example trajectory (black), and the momentary movement direction (red) and cone (blue) at four different points along the trajectory. Grey circles show (from bottom to top) starting sphere, via-sphere and target sphere. At each point along the trajectory, the cone encompasses all possible directions aimed anywhere at the target. **B:** Difference between the momentary direction and the closest cone surface as function of distance-to-targets. **C:** Speed as function of distance-to-targets. Samecolored markers in A-C refer to the same timepoints during the movement. The POC is defined as the point at which the difference between current direction and cone surface starts to decrease monotonically (purple marker) until in-cone (brown marker), or, if applicable, at a local speed minimum (orange marker) within the epoch defined by the purple and brown markers.

By defining the POC^cone^ like this, the commitment to the target is not necessarily assigned to the point in time when the effector direction for the first time changes into the general direction of the target (e.g. left of the vertical midline if the goal is located somewhere in the left of the workspace). Instead, the POC^cone^ is assigned to the point when the effector direction starts changing towards the surface area of the cone and keeps pace with the cone’s ever-changing orientation to finally enter the cone. The orientation and the opening angle of the cone, which define the range of directions aimed at the target, dynamically change because they are position dependent. For example, with smaller distance from the target the opening angle increases. As long as the current direction stays within the cone after entering it, the movement is considered to be “on target”. This means, once inside the cone, increases in the deviation of the current direction from the direction aiming at the center of the target are allowed, since the methods acknowledges the finite size of the target. Thus, the cone method takes into account that a larger target as well as decreasing distance from the target reduces demands on accuracy of the movement direction.

The cone method can be applied to both, 2D and 3D trajectories. Since in our experiments subjects produced 3D trajectories, we used the 2D projection onto the plane defined by the lateral deviation and distance-from-start vectors to demonstrate the cone method’s capability to handle 2D data. We computed the current direction at the trajectory’s sampling point k as the vector between the cursor position between k and k+1 (= k + 5ms since the trajectories were downsampled to 200Hz), and the closest in-cone direction as the tangent from the position of sampling point k to a d = 30mm circle placed on the target position. To apply the cone method to the original 3D trajectories, we again extracted the cursor position at k and k+1, and the target center’s position, all in XYZ coordinates. Since three positions in 3D space describe a plane, we were again able to compute both, the current direction and the closest in-cone-direction (i.e. intersection of the cone’s surface with this plane) in 2D as described above. Since this plane is position-dependent it is computed separately for each sampling point. For Experiment 1, we report the results of applying the cone method to 3D data (see Supplementary Information 1 for 2D results). For Experiment 2, we report the results of applying the cone method to 2D data, thereby disregarding movement components orthogonal to the task plane, since these orthogonal components did not carry any additional decision-related information in this factual 2D task.

We implemented three additional steps to the basic cone method to improve the POC^cone^ estimates. We adjust the POC^cone^ for two classes of cases where subjects temporarily moved out of the cone again, possibly without intent, after having already aimed the movement at the target, and we introduced a speed criterion.

First, we control for accidental out-of-cone slips by applying a 3° tolerance window to the cone (*tolerance*). This tolerance window comes into effect once the movement direction had already been inside the cone once. This means, when the movement direction moves out of the cone again by less or equal than 3°, we treat this epoch as “within-cone”. Second, we control for overshooting, i.e. moving inside the cone, followed by temporarily moving out again at the opposite side by more than 3° (*overshoot*). In Experiment 1, the opposite target was defined by the target direction plus 180 deg and identical nominal adjustment angle. If the movement direction moved away from the opposite target’s cone at a steeper rate than it moved away from the selected target’s cone, we defined this temporary slipping-out-of-cone as overshoot and treated the current direction during this epoch as “within-cone”. We applied this definition to both, 2D and 3D trajectories. Third, we acknowledge the possibility that the movement direction already by chance may have started to approach the cone, but subjects did not actually commit to the target yet. We assumed that in these cases, the actual commitment to the target later along the trajectory may be marked by the onset of an additional acceleration period (*speed*). Between the start of the movement adjustment (Figure 2, purple marker) and the point from which on the current direction remained in the cone (brown marker), we therefore searched for local minima in movement speed (Figure 2C). If a speed minimum was found, the POC^cone^ was defined as the location on the trajectory corresponding to this minimum (Figure 2, orange marker), otherwise the POC^cone^ remained as defined before (Figure 2, purple marker).

The three adjustments were applied hierarchically in the order they are presented above, i.e. the overshoot adjustment was applied to the data that resulted from applying the tolerance window, and the speed criterion was applied to the data that resulted from applying the overshoot control. All adjustments were applied to the data from both experiments, but their parameters were only tuned to Experiment 1. This is because only in Experiment 1 we were able to quantify the effects of the adjustments on the cone method’s performance, i.e. how they affected the proportion of POC^cone^ correctly recovered inside the via-sphere (see Supplementary Information 1 for a comparison of the different cone method adjustments). In the main text, for Experiment 1, we present the POC^cone^ that are estimated by the method including all three adjustments and being applied to the original three-dimensional trajectories, since this version of the cone method outperformed all other tested versions (see Supplementary Information 1 for an in-depth comparison). For Experiment 2, we only applied the cone method with all three adjustments. For brevity, we refer to this version as the “cone method”, as opposed to the “raw cone method” in Supplementary Information 1, which refers to the cone method without any of the three adjustments.

#### Cluster-based permutation test

Previous methods for quantifying choice latencies from trajectory data focused on group-level comparisons, i.e. comparisons between task conditions considering all trials in each condition. Therefore, in order to create a reference measure against which we can compare our novel approach as a control, we analyzed the data at the group level with an alternative method. While different specialized ad-hoc methods have been proposed for trajectory data in spatial selection tasks (e.g. Chapman et al., 2015), we here compare to a sensitive, yet rather general-purpose model-free method based on a cluster-based permutation test (*CP test*; Maris & Oostenveld, 2007; implementation used in this study based on Dann, Michaels, Schaffelhofer, & Scherberger, 2016). CP tests provide a means for non-parametric statistical comparison of time series data that accounts for the multiple comparison problem when performing statistical tests across many time points.

Here, we statistically determined when the trajectories towards two opposite targets started to branch. We tested per subject from which point onwards the lateral deviation between pairwise groups of trajectories started to be significantly different from one another using t-tests embedded in the CP test. For Experiment 1, we tested trajectories pooled across the upper two target positions versus trajectories pooled across the lower two target positions. We conducted separate CP tests per nominal adjustment angle. In Experiment 2, we tested trajectories towards the left versus the right target. We tested per subject, per SOA, and separately for gain and loss trials (see Supplementary Information 2 for a detailed account of how we applied the CP test). We treated the obtained significance onsets as point of overt commitment as recovered by CP tests (POC^CP^).

#### Statistical assessment of the POC^cone^ and POC^CP^ results

We analyzed the POC^cone^ and POC^CP^ results extracted from the trajectories in Experiment 1 and 2 using generalized linear mixed effects models (GLME, Matlab function **`fitglme`**). Whenever we fitted a model to proportions, we assumed binomially distributed data and used a logit link function. For each model, we started off by only including the main factors and added interaction terms where applicable. In these cases, we decided whether to include the interaction terms based on Akaike’s information criterion (AIC). Main effects are always reported from the models without interaction. Each model included, per subject, random intercepts and random slopes for each fixed effect (omitted in the model descriptions using Wilcoxon notation below).

In Experiment 1, first, we assessed whether the proportions of POC^cone^ recovered inside the via-sphere, prior to entering the via-sphere (*too-early-POC*), and after leaving the via-sphere (*too-late-POC*) depended on the steepness of the adjustment angle. We fitted the following GLME separately on each of these three proportions.

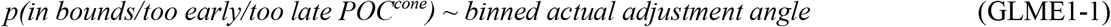

The actual angle about which a movement had to be adjusted inside the via-sphere deviated from the nominal adjustment angle defined by the spatial stimulus arrangement. This is because of the finite size of the starting-, via-, and target spheres, as well as the 30° range for the movement direction inside the movement corridors. We therefore estimated the actual adjustment angle (see Supplementary Information 3) for use in this (GLME1-1) and the following analyses. We discretized the resulting adjustment angle estimates in 10° wide bins starting at 0°. Trials with actual adjustment angles smaller than 10° and equal to or larger than 80° were omitted from all GLMEs that included these binned actual adjustment angles because the respective bins were made up of only 1-6 trials for certain subjects (all other bins contained at least 9 and on average 27 trials per subject).

Second, we assessed whether the POC^cone^ itself varied as function of the steepness of the adjustment angle. To measure the effect of adjustment angle on POC^cone^ independently of the distance between starting- and via-sphere, we used the POC^cone^ relative to the via-sphere entry point:

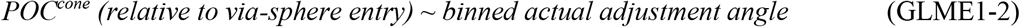

Third, we quantified how well POC^cone^ and POC^CP^ matched depending on the adjustment angle. For this, we grouped the trials according to their nominal adjustment angle instead of their actual adjustment angle since the former had to be used to group the data for the CP tests. We performed the following GLME on the POC^CP^ and, to directly compare between the two, also on the POC^cone^ data:

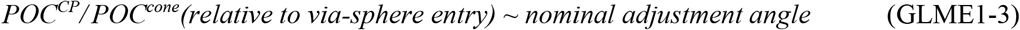

In Experiment 2, we converted the POCs obtained by both, the cone method and the CP test, into their corresponding movement time stamps (*time of overt commitment*, TOC), to directly relate them to the value cue SOA. First, we used the results of the cone method to group trials into three partially overlapping classes to demonstrate how we can identify trials with pre-movement commitment (TOC at movement start), potential guesses (TOC prior to value cue onset + 50 ms), and the combination of the two (“all early”; i.e. all trials without putative online decisions that follow value cue onset, regardless of whether this was the result of a pre-movement commitment or a commitment prior to value cue onset) respectively. We separately fitted these proportions as function of SOA to study how they depended on the timepoint at which the informed-choice-enabling value cue became available:

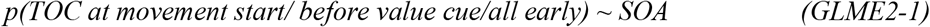

Second, we assessed how SOA and gain vs loss frame influenced the point of overt commitment. We computed the following GLMEs, only including trials in which subjects chose the higher valued option:

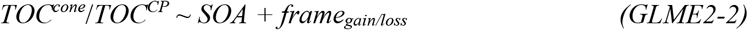

In the results section we only report significant effects of each GLME. The full results of each model can be found in the corresponding tables in Supplemental Material 4.

## Results

### Experiment 1

In Experiment 1, we assessed the cone method’s capability to correctly recover POCs inside the via-sphere as proxy of how close the POC^cone^ estimates are to the true point of overt commitment trial-by-trial. We further assessed how the cone method performed in comparison to a CP test in across-trial data.

#### Rate of successful in-bounds POC^cone^ estimates

Figure 3A shows how the majority of all POC^cone^ were detected within the via-sphere across all three possible via-sphere distances from the starting position. Except for putative outliers, the distributions of POCs were narrower than the via-sphere diameter and approximately centered on the via-spheres. This shows that the cone method produced plausible POC estimates, nicely tracking the position of the via-sphere.

**Figure 3.**
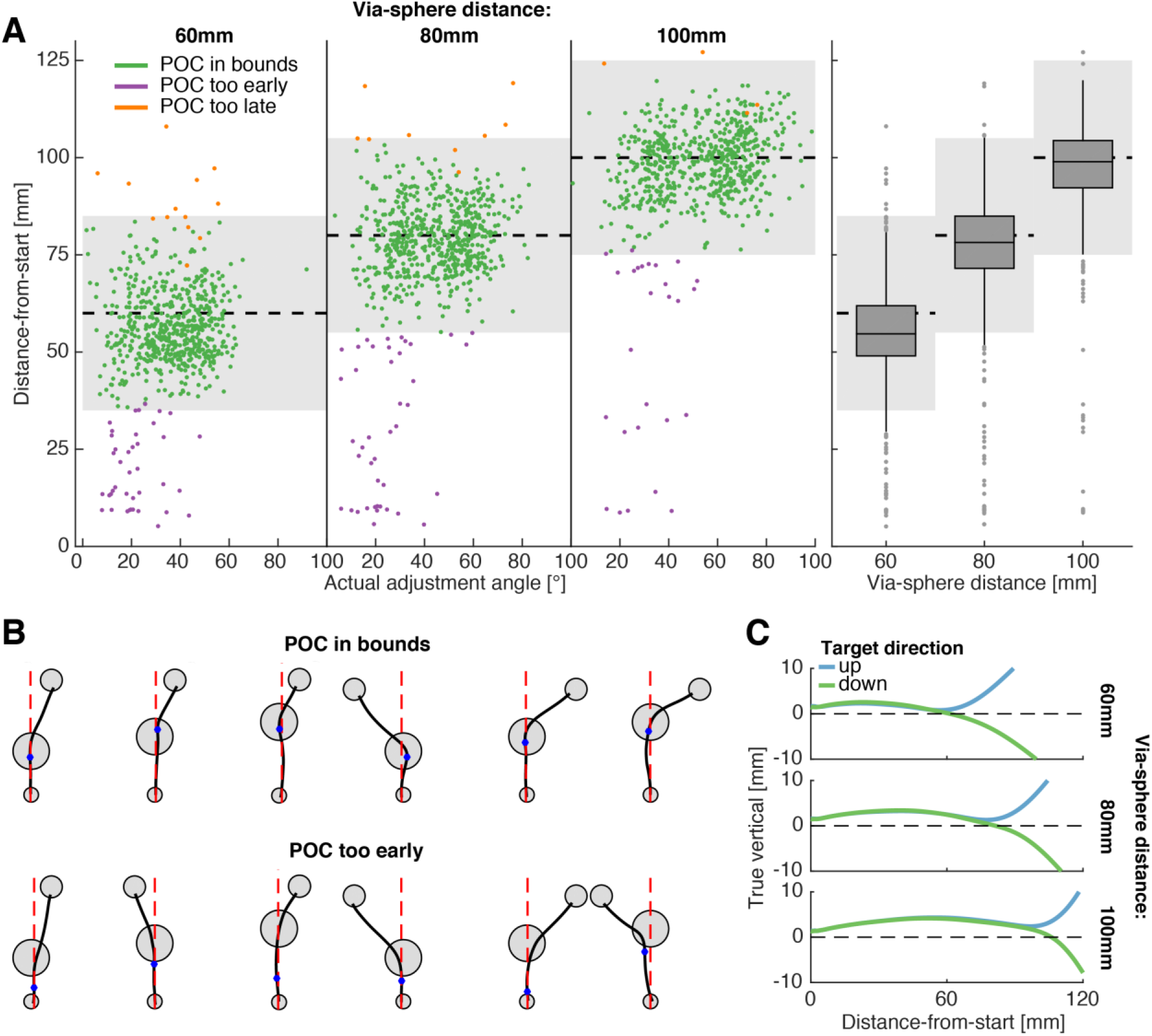
POC^cone^ estimates and early movement biases that lead to too-early POC^cone^. **A.** POC^cone^ of all subjects for each of the three starting-sphere to via-sphere distances as function of the actually produced directional adjustment angle of the trajectory (left three panels). The gray-shaded areas indicate position (dashed horizontal line) and maximum diameter of the via-sphere. POC^cone^ classified as out-of-bounds that appear within the gray area are due to the subject entering/leaving the 3D via-sphere above/below its maximum diameter. The right panel shows the same data, pooled across adjustment angles, including a box-and-whisker plot representing median and 25^th^/75^th^ percentiles, and the 1.5-times interquartile ranges. **B.** Example trials (2D projections of the trajectories, as in Figure 1D) and POC^cone^ estimates (blue dots) detected in-bounds (upper row) or too early (lower). Dashed red lines indicate the distance-to-targets axis. **C.** Average trajectories across all trials per via-sphere distance in the side view (manipulator workspace-defined X- and Y-axes). The Y-axis is magnified threefold to emphasize the degree of early movement bias due to ballistic lifting of the hand during anterior transport.

However, the cone method also produced a certain fraction of too-early-POCs, especially (and not surprisingly) at low adjustment angles. These putative misclassifications are instructive for understanding the limitations of the method. In cases of POC detection prior to entering the via-sphere, example trajectories often either show a smooth, shallow adjustment towards the target position, with the absence of a clear turn in the trajectory (Figure 3B, lower row from left to right: example trials 1-5), or a movement in the general direction of the target sphere prior to its onset and the subsequent within-via-sphere turn (example trial 6). We assessed whether movement biases prior to entering the via-sphere could have accounted for the higher number of too-early POCs at low adjustment angles. Since there were tolerances in movement curvature during the outside-via-sphere movement corridors, such biases were possible. Figure 3C shows how subjects indeed on average moved towards the viasphere in an upward ark, entering the via-sphere from slightly above. In the case of downward targets, this led to smaller actual adjustment angles (compared to the actual adjustment angles towards upper targets within the same nominal adjustment angle condition; see Supplementary Information 3). Consequently, the movement direction upon target onset already coincided with the general target direction. In summary, the movement constraints outside the via-sphere did not fully prevent the subjects from making slightly curved movements towards the via-sphere. This led to partial misclassifications which should not be attributed to a suboptimal performance of the cone method, but rather reflect limited compliance in behavior (see Supplementary Information 3 for a discussion on the performance of the cone method at small adjustment angles independent of the susceptibility to initial movement biases).

#### POC estimates as function of adjustment angle

We assessed quantitatively how the steepness of the actual adjustment angle influenced the number of out-of-bounds POCs. For this, we separately fitted GLME1-1 to the proportions of in-bounds/too-early/too-late POC^cone^, respectively (Figure 4A). In-bounds POCs on average accounted for 93% of all trials. Their proportion relative to the total number of trials within each adjustment angle bin increased with increasing adjustment angle (*log odds ratio* = 0.061; *p* < .001) and stayed above 95% for angles equal or larger than 30 deg. This increase was almost exclusively attributable to the decrease of too-early POCs (total average = 6%) with increasing adjustment angle (*log odds ratio* = −0.084; *p* < .001), in line with the pattern visible in the raw data presented above. Too-late-POCs only accounted for a total average of 1% of all trials and did not significantly vary with adjustment angle (*p* = .79). We did not perform analyses on the proportions of out-of-bound POC^CP^ as all but one (too-early-POC^CP^ at 25.76°) out of the 48 (total across subjects) CP tests yielded in-bounds POCs.

**Figure 4.**
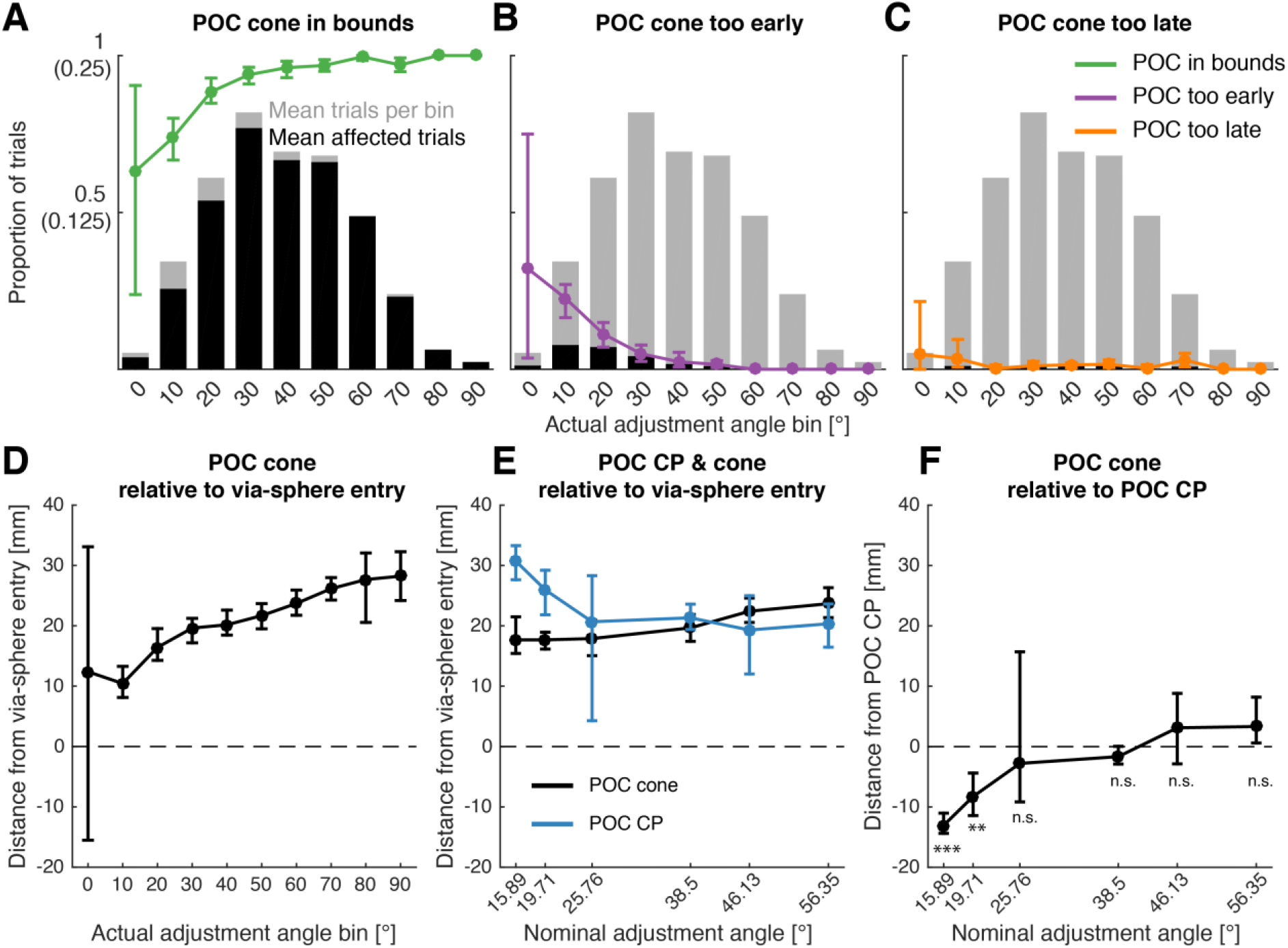
Likelihood and position of in-bound POC localization. Top row: The line graphs show the average per-subject proportions of POC^cone^ recovered in bounds (**A**), too early (**B**), and too late (**C**) as function of actual adjustment angle. The gray bar graphs show the average per-subject proportion of trials in each bin, out of the 240 trials per subject across all bins. Note, this data is identical in A-C. The black bar graphs show the corresponding proportions of in-bounds, too-early-, and too-late-POC^cone^, respectively. The Y-axis tick marks without parentheses refer to the line graphs, the Y-axis tick marks in parentheses refer to the bar graphs. Bottom row: Across-subject average of the within-subject average POC^cone^, relative to the via-sphere entry-position and as function of the actual adjustment angle (**D**), POC^cone^ and POC^CP^ relative to the via-sphere entry point and as function of the nominal adjustment angle (**E**; data averaging as in D), and POC^cone^ relative to POC^CP^ and as function of the nominal adjustment angle (**F**; data averaging as in D). Asterisks in D are one-sample t-test p-values with ** = *p*<0.01, *** = *p*<0.001, and n.s. = not significant. All error bars are bootstrapped (N = 2,000) 95% confidence intervals of the mean.

Figure 4B shows the POC^cone^ and POC^CP^ relative to the via-sphere entry point and relative to each other. POC^cone^ were recovered further up along the distance-from-start axis with increasing actual adjustment angle (left panel; β = 0.228, *p* < .001; GLME1-2). This pattern was weaker, but still observable, when grouping the POC^cone^ according to the nominal adjustment angle (middle panel; β = 0.16, *p* < .001; GLME1-3). In comparison, the POC^CP^ were recovered earlier with increasing nominal adjustment angle (middle panel; β = −0.217, *p* = 0.012; GLME1-3). We performed one-sample t-tests on the difference between POC^cone^ and POC^CP^ per nominal adjustment angle to assess at which adjustment angles the two POC measurements yield different results (right panel). Due to the positive slope of POC^cone^ and the negative slope of POC^CP^, both methods recovered different POCs at small adjustment angles (15.89° & 19.71°: *t*(7) = −14.21, *p* < .001 & *t*(7) = −4.43, *p* = 0.003; the significance of all t-tests was determined at the Bonferroni-corrected alpha level of 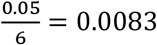). POC^cone^ and POC^CP^ were similar at nominal adjustment angles of 25 degrees and larger (all *p* > .08).

### Experiment 2

Subjects performed a go-before-you-know decision-making task to targets that were associated with different monetary outcomes. The outcomes were cued at different SOAs relative to the go-cue. We used the cone method to classify trials in pre-movement and peri-movement decisions and compared how well the cone method and the CP test were able to capture the known and expected effects of value cue SOA and gain- versus loss frame on the times of overt commitment.

#### Single-trial trajectory classification using the cone method

Figure 5A shows all trajectories of a single example subject, grouped by the experimentally controlled SOA (columns). Additionally, we classified the trials based on whether the cone method estimated the TOC to have occurred during the movement (top panels) or already at movement start (bottom panels). Visual inspection shows how the classification of trajectories based on the cone method successfully split the data into movements with at least one clear turn and thus putative “online” commitment to choice (top) versus direct-to-target movements (bottom). A few trajectories in the bottom panel deviated from this pattern but showed a single smoothly arched movement towards the chosen target (e.g. 283ms SOA subpanel).

**Figure 5.**
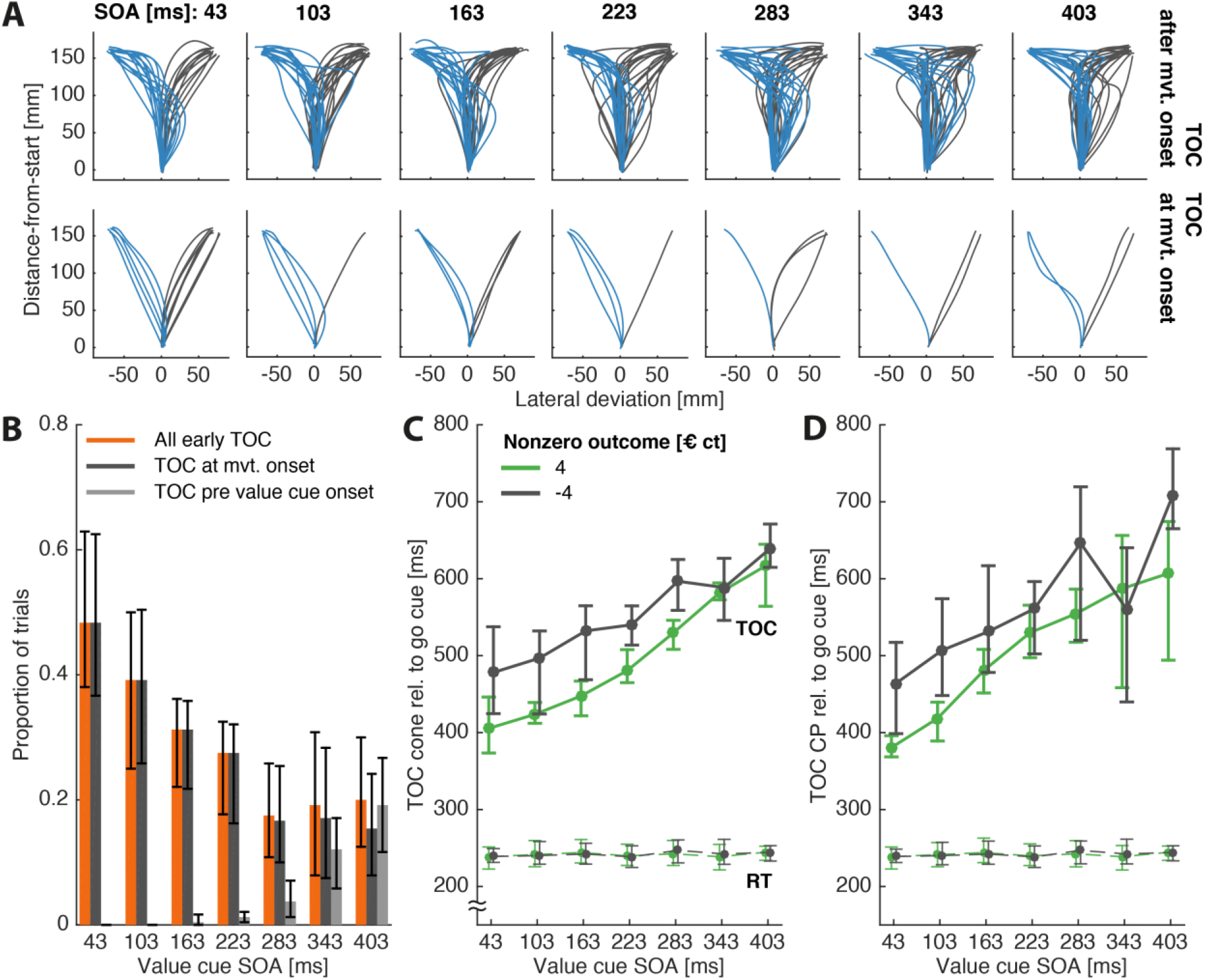
Experiment 2 results. **A.** Single subject example trajectories (all successful trials) split by whether TOC^cone^ was estimated to have occurred during the movement (top) or directly at movement start (bottom). **B.** Across subject mean proportions of TOC^cone^ at movement start, before value cue onset + 50ms, and the total of the two. Note that the proportion of all early TOC does not need to be the sum of the other two proportions since a single TOC can be both, estimated at movement start, and estimated prior to value cue onset. **C.** TOC^cone^ (solid lines) and movement initiation times (RT, dashed lines) relative to the go-cue (higher-value choices only). **D.** TOC^CP^, same conventions as in C. All error bars are N = 2,000 bootstrapped confidence intervals of the mean.

When designing a go-before-you-know experiment to measure TOCs/POCs, it is important to select an appropriate SOA for the stimulus that determines target selection (here: value cue). If this stimulus is displayed to early, subjects can select the target prior to movement start. If it is displayed to late, subjects might resort to guessing because they run out of time to adjust their movement in response to the stimulus. In both cases, the resulting trajectories are uninformative regarding the TOC in response to the stimulus. We therefore demonstrate, using the cone method, how to determine an optimal SOA that yields the highest number of informative trajectories. We sorted the trajectories into “TOC at movement start” (value cue displayed too early), “TOC prior to value cue onset” +50ms (value cue displayed too late), and “All early TOC” (POC either at movement start or before value cue onset; Figure 5B). The proportion of TOC at movement start decreased with increasing value cue SOA (*log odds ratio* = - 0.006, *p* = .002; GLME2-1), while the proportion of TOC before value cue onset increased with increasing SOA (*log odds ratio* = 0.0161, *p* < .001; GLME2-1). Both was to be expected, given the explanation above. Critically, the proportion of “All early TOC” overall decreased with increasing SOA (*log odds ratio* = 0.005, *p* = 0.005; GLME2-1), and was lowest at 283ms, which made it the optimal value cue onset latency relative to the go cue.

#### Effects of value cue SOA and gain- versus loss frame on time-points of overt commitment

We assessed how gain/loss framing and value cue SOA influenced the time subjects needed to commit to the higher-value target (Figure 5C), similarly to Chapman et al. (2015). We estimated the TOC^cone^ using the cone method and fitted GLME2-2 to the resulting TOCs, only including trials where subjects chose the higher-value target and pooling across higher-value target location (left versus right). TOCs were on average faster by 55ms (β = −55.215, *p* < .001) in gain trials compared to loss trials. Further, TOCs increased on average at 52.5% the rate of the increasing SOA (β = 0.525, *p* < .001). This increase existed in both, gain- and loss trials, but was larger for gain- compared to loss-trials (interaction SOA*gain-/loss frame β = 0.179, *p* = 0.003). This difference in slope was accounted for by the gain- versus loss effect being most prominent at low to mid SOAs and almost absent at high SOAs.

We additionally estimated the TOC^CP^, i.e. the timepoint underlying the position at which trajectories towards the left and right target began to differ significantly from one another. We obtained separate TOC^CP^ per subject, SOA, and gain- versus loss frame. We were unable to detect a significant separation of the two groups of trajectories between high- and low-value targets for 5 out of the 84 computed CP tests, due to the trajectories exhibiting high within-group variability. We compared the results of the remaining CP tests with the cone method by fitting GLME2-2 to the corresponding TOC^CP^ estimates (Figure 5D) and obtained comparable results (main effect of SOA: β = 0.608, *p* < .001; main effect of gain-/loss frame *β* = 64.375, *p* = .001).

## Discussion

We developed and tested the cone method to estimate on a single trial basis when online commitment to move towards a spatial target becomes detectible in ongoing movements. The cone method estimates these points of overt commitment (POC) by identifying the point where the momentary movement direction starts to become adjusted monotonically towards the range of possible movement directions aimed at the target, and additionally factors in the movement speed as marker for commitment to further improve this estimate. We conducted two experiments. In Experiment 1, subjects performed instructed go-before-you-know reaches to one of eight potential targets, the position of which was revealed only shortly before entering a via-sphere. Only during the movement portion inside the via-sphere subjects could curve their movements towards the target position, thus establishing a “ground truth” data set with a bounded range of commitment points based on which we established a proof-of-concept and fine-tuned the cone method. In Experiment 2, subjects performed unconstrained go-before-you-know reaches to one of two potential targets with varied monetary outcomes. We were able to estimate POCs in this choice task on a trial-by-trial basis, confirming known dependencies of choice latency on cue timing and reward framing. We also demonstrated the cone method’s ability to classify trials as direct reaches without online decision versus curved reaches with online decisions.

### Advantages of Measuring Online Decision Processes Over the Use of Reaction Time Paradigms

Conventional laboratory settings usually impose a strict and temporal segmentation of stimulus presentation and behavioral readout. Stimuli are presented before subjects are able to initiate their response and discrete behavioral readouts such as button presses prevent subjects to revise their response once initiated. Only under rare circumstances however, one needs to withhold all movement until a trigger allows a sudden response, the latency of which can be measured as reaction time. Natural behavior instead unfolds as result of a continuous sensorimotor integration process (Pezzulo & Cisek, 2016). In ecologically more valid situations, action selection and movement control are parallel processes. These processes are characterized by similar minimization principles (Morel et al., 2017; Shadmehr et al., 2016) and movement paths can be selected rapidly in response to sudden perturbations of ongoing movements (Nashed et al., 2012, 2014; see also Gallivan et al., 2018; Wispinski et al., 2020 for review), underscoring the link between selection and control of action.

Novel experimental approaches seek to break up artificial temporal constraints in favor of more naturalistic behavioral paradigms in which sensory encoding and action selection can take place during continuously evolving behavior. Go-before-you-know tasks (e.g. Gallivan & Chapman, 2014), including mouse tracking tasks (e.g. Freeman et al., 2011), maintain a fixed trial structure but enable subjects to start their action during continuing stimulus-guided action selection. Other paradigms allow subjects to sequentially make multiple decisions within a given trial while continuously moving around the task environment (Diamond et al., 2017; Michalski et al., 2020), effectively abolishing the one-decision-per-trial structure for more naturalistic choice sequences. Also, coordination of action with others, be it in a cooperative or a competitive setting, requires continuous integration of selection and control when the opponents’ actions are mutually visible and leads to specific dyadic choice behavior, like leader-follower strategies in transparent games (Möller et al., 2020; Unakafov et al., 2020).

In summary, decision-making and action execution are highly linked, rather than the latter only being a mere behavioral confirmation of an already finished, preceding choice process. Estimating decision processes during continuously evolving action will thus be an important task for studying increasingly ecologically relevant behaviors.

### Methods to Analyze Movement Trajectories as Marker of Online Decision Processes in Spatial Selection Tasks

The paradigms described above capitalize on our ability to online-select and revise movement targets by using movement trajectories to study cognitive processes such as stimulus processing and decisionmaking. They provide richer data than conventional “decide, then act” reaction time paradigms. In turn, they lack an inherent measure for the duration of the decision process. In go-before-you-know tasks, reaction times are often uninformative (Gallivan & Chapman, 2014), while in paradigms without segmented trial structure reaction times cannot be measured as subjects move uninterruptedly between their consecutive decisions. A variety of methods, especially for go-before-you-know paradigms, has been established to investigate the complexity and temporal structure of decision processes during ongoing movements. Here, we discuss two classes of such methods and how the cone method complements them or provides an advantage over them. The cone method is the first method to our knowledge that allows to directly quantify decision times on a single trial basis, thereby combining the advantages of investigating decision processes during ongoing movements and conventional reaction time paradigms. Additionally, the cone method can also be applied to 3D trajectories, which is not easily obtained with the methods described below. Therefore, it provides a distinct addition to existing tools used to study decision-making during ongoing movements.

#### Summary measures

Measures such as maximum deviation (Freeman et al., 2011) or area under the curve (Freeman & Ambady, 2010) per each trajectory provide a summary value of how close this trajectory is to an idealized straight-to-target movement. What is typically measured is the size of the maximum deviation of the trajectory orthogonal to a straight line from movement starting point to endpoint or the area between the trajectory and said line, respectively. According to this logic, a trajectory following the straight line between movements starting point and target marks the extreme of a decision prior to movement start. These summary measures, which translate the course of the trajectory relative to an idealized reference trajectory into a single scalar number, allow ordinal statements about the relative decision latency between trajectories. The smaller the maximum deviation or area and hence the higher the proximity to a straight-to-target movement, the earlier the decision.

In an Experiment similar to our Experiment 2, Chapman and colleagues (2015) translated area measures into decision time differences between two experimental conditions by estimating how much earlier the value cue had to be presented in loss-compared to gain trials to evoke trajectories with similarly sized areas. They found that the value cue had to be presented on average 100ms earlier in loss trials compared to gain trials, which means deciding against a loss target took subjects 100ms longer than deciding for a gain target. When using this method, the successful estimation of decision times depends on how reliably the trajectory areas change with increasing SOA. Without such SOA-dependency of the decision time, the approach of varying SOA for identifying similar-size trajectories between two experimental conditions does not work. In our Experiment 2, the area measures decreased only very little with increasing SOA, while at the same time the effect of gain-versus loss trials on the areas was large. As consequence, the approach of Chapman and colleagues would not have allowed us to determine the amount of SOA difference between gain and loss trials that would equal the area measures between these two conditions (data not shown).

When we instead measured the difference in TOC with both, the cone method and the CP test, a ~50ms TOC difference between gain and loss trials and a ~50% increase in TOC relative to the increase in SOA became evident, suggesting that these measures are more sensitive than summary measures. With the cone method, we were able to measure the differences between gain- and loss trials directly and without having to resort to trajectory areas and their dependency on value cue SOA.

Also, cognitive dynamics like changes of mind easily will be underappreciated with summary measures since they lead to trajectories of more complex shape. Two trajectories with identical time-point of final commitment will differ in their areas or deviation measure, depending on the history of preliminary choices that might have preceded the final commitment, e.g. because they curved the trajectory in S-shape. The cone method is designed to identify the final point of commitment leading to the ultimate selection.

Finally, methods depending on a reference trajectory against which an area or deviation is measured (Freeman et al., 2011; Freeman & Ambady, 2010), or which determine areas otherwise (Chapmann et al. 2015), require a defined starting and end point, between which the measure is taken. The cone method can be applied to ongoing movements without defined starting point, as long as the target region is defined.

#### Time series analyses

In contrast to the summary measures described above, time series analysis such as per-time-point regression (Dotan et al., 2019; Scherbaum et al., 2010) allow to measure the temporal evolution of the decision making process. Separate linear regression models are fitted at each sampling point of all trajectories. The decision-enabling variables (e.g. the value cue from our Experiment 2) are the predictors in these regression models. The coordinates of the trajectories along the dimension defined by the axis between the potential movement targets (the lateral deviation in our experiments) are the dependent variable. The resulting curve of regression weights quantifies the size and onset of the predictors’ effects on the trajectories and thus allow to determine when these variables start to influence the decision process. The regression weights do not tell when this process is finished, though. The cone method here offers a complementary metric. The POC estimates when the decision process is completed.

A method to directly infer POCs from movement trajectories is to determine when trajectories towards spatially separated targets start to branch, as we statistically assessed using a cluster-based permutation test in our Experiments 1 and 2. Effectively, this is a differential measure, with the consequence that the POC estimate depends on how the trajectories towards either target are shaped in relation to each other. This comparison of trajectories leads to two drawbacks that we were able to circumvent with the cone method, which estimates POCs independently based on a single trajectory.

Firstly, one needs to assure that the trajectories to the alternative targets do not branch prior to the putative POCs, as would be the case, for example, if differently high instruction probabilities cause subjects to bias their movements towards the more likely target prior to the instruction onset. Using methods like the CP test could then cause erroneously early POC estimates. With the cone method, instead, a pre-commitment separation of trajectories does not lead to premature POC^cone^ estimates as long as the actual commitment is still marked by a clear turn in the trajectory. This can be facilitated via a sufficiently large target separation angle, as demonstrated by the dependency of the cone method’s performance on the adjustment angle in our Experiment 1.

Secondly, a large variability within the groups of trajectories towards either target (e.g. due to a large variability in the true POCs), especially in conjunction with a small group separability (e.g. due to spatially close targets), cannot be properly captured by trial-averaging approaches. If the true POCs are widely distributed along the distance to the target array, this can lead to erroneously late or even no POC estimates, i.e. only once most of the trajectories are adjusted to the targets and the within-group variability does not overlap anymore. This means, group-level trajectory comparisons are biased towards late POCs when decision times are more variable in time. The late POC^CP^ estimates (relative to the POC^cone)^ at small nominal adjustment angles resulting from spatially close targets in our Experiment 1 are an example for this effect. The cone method instead captures this variability and allows to extract the unbiased distribution of decision times. We were mostly able to control for differences in the variability of the decision times with the strict movement constraints in our Experiment 1. Nevertheless, we measured later POC^CP^ at small, compared to larger lateral target offsets. In proper choice experiments, however, it is not meaningful to constrain the time of movement curvature as it is the main outcome variable of interest. In our Experiment 2, the results of the CP test were similar to the results of the cone method, but we were unable to obtain significant trajectory differences for 5 out of 84 CP tests, indicating limited sensitivity of the CP test. The cone method instead was able to properly capture the mean and variability of the POCs. As a single-trajectory measure, the cone method aggregates the within-group data after estimating the commitment points per each trajectory, as opposed to group-level measures doing the opposite, allowing to estimate the distribution of commitment points.

### Limitations and Recommendations when Applying the Cone Method

In Experiment 1, we found a 25° lateral target offset (equivalent to the 50° target separation in Experiment 2) to produce sufficiently high trajectory adjustment angles for the cone method to validly estimate the POC^cone^. We consider these estimates valid since in this range of larger adjustment angles the fraction of in-bounds recoveries was close to 100% and since the average POC^cone^ across trials and POC^CP^ estimates were highly similar. Consequently, we find it reasonable to argue that both methods produced plausible POC estimates at mid to high adjustment angles, only that the POC^cone^ achieves this also at single-trial basis, not just on average. Assuming that the true point of commitment on average does not vary with nominal adjustment angles, the fact that the POC^cone^ shows a flat curve towards low adjustment angles (Figure 4B middle) suggests a more truthful POC^cone^ estimate within this range of low adjustment angles, compared to the POC^CP^, which seems biased towards later POCs. Yet, also POC^cone^ suffers from small adjustment angles, as the fraction of too-early recoveries of POC was higher for small angles (Figure 4A).

### Conclusions

We developed the cone method as algorithm to estimate points of overt commitment from go-before-you-know trajectories on a single trial basis. The method is applicable to both, two- and threedimensional movements. We demonstrated how the cone method can be used as reaction time analog for online decisions during movements and to classify those movements based on the decision time relative to specific task events. Even though we validated the cone method using three-dimensional manipulator-based reaching movements, we believe it can also be applied to other settings where subjects make decisions midflight during ongoing, tracked movements such as tap-to-touchscreen paradigms (e.g. Gallivan & Chapman, 2014) and mouse tracking (e.g. Freeman et al., 2011; Spivey et al., 2005). We consider the method also suited to analyze decision processes in free continuous movements during naturalistic tasks like chasing pray or foraging, when both, trial-averaging methods and 2D-constrained trajectories are not applicable.

## Data Availability

All Matlab scripts required to perform the analyses and output the figures described/presented in this study, and the underlying data are available on https://github.com/PhilUlb/conemethod

## Acknowledgements

We thank Thérèse Morgenstern for help with data collection for Experiment 2. This work was supported by the German Research Foundation in the context of the Collaborative Research Center “Cellular mechanisms of sensory processing” (DFG, grant SFB 889 C4 to A.G.) and by the European Commission in the context of the Plan4Act consortium (H2020-FETPROACT-16 732266 WP1 to A.G.).

