## Supplementary Information for "The Cone Method: Inferring Decision Times from Single-Trial 3D Movement Trajectories in Choice Behavior"

### Supplementary Information 1: Performance gain by adding extra criteria to the raw cone method (Experiment 1)

In the main text, we described the results of the cone method after the addition of three adjustments that improved the method's capability to recover the POC$^{cone}$ inside the via-sphere. We hierarchically added: a tolerance window to the cone surface (Supplementary Figure 1-1 & 1-2: "tolerance"); an adjustment to exclude out-of-cone-slips due to directional adjustments of the movement that exceeded the lateral offset of the target ("overshoot"); and a speed criterion that shifted the POC$^{cone}$ to acceleration onsets that occurred between the beginning of the movement adjustment and entering the cone ("speed"; see main text for a more detailed description). Here, we compare the proportions of too-early-POC$^{cone}$, too-late-POC$^{cone}$ and in-bounds POC$^{cone}$ between each of these adjustments. Additionally, we compared these values between the cone method including all adjustments applied to the original, three-dimensional trajectories and their projections on the lateral deviation – distance-from-start plane (Supplementary Figure 1-1 & 1-2: "2D").

#### Proportions of Trials Affected Due to Additions to the Cone Method

Supplementary Figure 1-1A shows the proportions of POC$^{cone}$ recovered inside the via-sphere, before entering the via-sphere, and after leaving the via-sphere, respectively, as function of the actual adjustment angle. Each version of the cone method contained the hierarchically lower adjustments, i.e. the data shown from the cone method version with overshoot adjustment also included the tolerance window around the cone surface and the data shown from the cone method version with speed criterion also contained both, the tolerance window and the overshoot adjustment. One can visually appreciate how for all cone method versions the proportion of too-early-POC$^{cone}$ was reduced with increasing actual adjustment angle. This effect was approximately equally sized in all four versions of the cone method applied to the 3D trajectories, and larger for the POCs obtained from the 2D projections. The distribution of too-late-POC$^{cone}$ across actual adjustment angles was flat and adding the overshoot adjustment eliminated almost all cases. Conversely, the proportion of POCs recovered within the via-sphere increased with increasing actual adjustment angle in all versions of the cone method and reached a plateau of almost 100% when the cone method was applied with all three additions on both, 3D and 2D trajectories.

Supplementary Figure 1-1B shows the same data as Supplementary Figure 1-1A, pooled across adjustment angle. We compared the proportions of POC$^{cone}$ in bounds, too-early, and too-late of the neighboring cone method versions as displayed in the figure, using Wilcoxon signed-rank tests at an alpha level of .025 (Bonferroni correction since each sample is used for up to two tests, depending on the number of neighboring samples). Applying the cone method with the speed criterion on the 3D trajectories yielded a higher proportion of POCs recovered in bounds, both in comparison to omitting the speed criterion and to applying the cone method with all adjustments to the 2D trajectories (both $p$ = .008). Both, adding the tolerance window and the overshoot adjustment, resulted in a higher proportion of POCs recovered too early, while adding the speed criterion decreased this proportion to the lowest value of all tested cone method variants applied to 3D trajectories, as well as in comparison to the cone method with speed criterion applied to 2D trajectories (all $p$ = .008). The proportion of POCs recovered too late decreased between the raw cone method and the cone method including the tolerance criterion ($p$ = .008), and between the cone method including the tolerance criterion and the cone method including the overshoot criterion ($p$ = .016). Adding the speed criterion did not significantly increase the proportion of too-late-POCs but applying the cone method with all additional criteria to 2D instead of 3D trajectories again decreased the proportion of too-late-POCs ($p$ = .008). In summary, applying the cone method with all three additions to the 3D trajectories yielded both, the highest proportion of POCs recovered in bounds (93%) and the lowest proportion of POCs recovered too early (6%), and a very low proportion of POCs recovered too late (1%; Supplementary Figure 1-1B).

#### Extent of POC Adjustment Due to Additions to the Cone Method

We additionally investigated the impact of each addition to the cone method on the POC estimate itself. Supplementary Figure 1-2 A&B show the proportion of $POC^{cone}$ estimates affected by each addition to the raw cone method. One can visually appreciate how adding the tolerance window and the overshoot adjustment affected the $POC^{cone}$ estimate in a small proportion of trials (mean = 7% and 5%, respectively), uniformly distributed across the different adjustment angles.

Adding the speed criterion affected on average 64% of trials, with fewer trials affected at low adjustment angles due to a mixture of two reasons. Firstly, the movement direction often was already in-cone prior to a speed minimum. Secondly, at small adjustment angles, the difference between movement direction and cone surface decreased in a less stable manner. Late (i.e. after the movement direction had already started to approach the cone’s surface) local maxima in this difference that determined the POC occurred after a speed minimum which excluded it from being considered by the speed criterion. To catch up with the cone’s surface, the current movement direction needs to approach the cone surface at a higher rate than the cone surface would move away from the current movement direction if the latter remained stable. At small adjustment angles (and consequently, a small rate at which the cone surface moves away), small fluctuations in the current movement direction may already violate this requirement, leading to local increments in the difference between movement direction and cone surface (which determined the POC) although the difference had generally already started to decrease until in-cone. At large adjustment angles, such small fluctuations only affected the slope of the movement direction – cone difference, without affecting the sign.

Applying the cone method with all additions to 2D instead of 3D trajectories changed the POC estimate in 29% of trials, with fewer trials affected at high adjustment angles. Similarly to what caused a subset of the low number of speed-criterion affected trials outlined above, late peaks in the difference between current movement direction and cone surface were more frequent in the 3D than in the 2D trajectories. This was to be expected since using the 2D projections eliminated one dimension of movement fluctuations that could have accounted for this effect. At large adjustment angles, using 3D versus 2D trajectories resulted in identical POC estimates for a large proportion of trials because again, these small fluctuations had no impact on the distribution of peaks in the movement direction – cone difference. Moreover, we used the speed curves obtained from the 3D trajectories in both cases, which resulted in identical POCs whenever the speed criterion was fulfilled.

Supplementary Figure 1-2 B&C show by how much the $\text{POC}^{\text{cone}}$ estimates were adjusted after applying the additions to the cone method. The tolerance window and overshoot adjustment had a large, highly variable effect. This comes unsurprising, as these “out-of-cone slips” could occur anywhere after the first within-cone portion of the movement and adjusting for them always placed the POC to the point where the current direction started to move towards the cone. The effect of adding the speed criterion was small. Under the assumption that a speed increase following target selection indicates commitment, this shows that the raw $\text{POC}^{\text{cone}}$ estimates were close to the true point of commitment, as they lead the adjusted $\text{POC}^{\text{cone}}$ estimates by approx. only 10mm. The influence of using 3D trajectories versus their 2D projection highly depended on the adjustment angle. Again, we attribute the larger difference at low adjustment angles to the large influence of small fluctuations in movement direction on the difference between movement direction and cone surface.

In summary, even though both, the number of affected trials and the amount of the POC adjustment varied substantially across the different additions of the cone method, the overall effect on the average POCs (including affected and unaffected trials) was small (Supplementary Figure 1-2D, “All trials”). However, the large effect of small fluctuations in movement direction on the POC estimates, as demonstrated by the proportions of trials affected by adding the speed criterion/using 2D instead of 3D trajectories and the magnitude of POC change between 3D and 2D trajectories, indicate a less stable performance of the cone method at small adjustment angles.

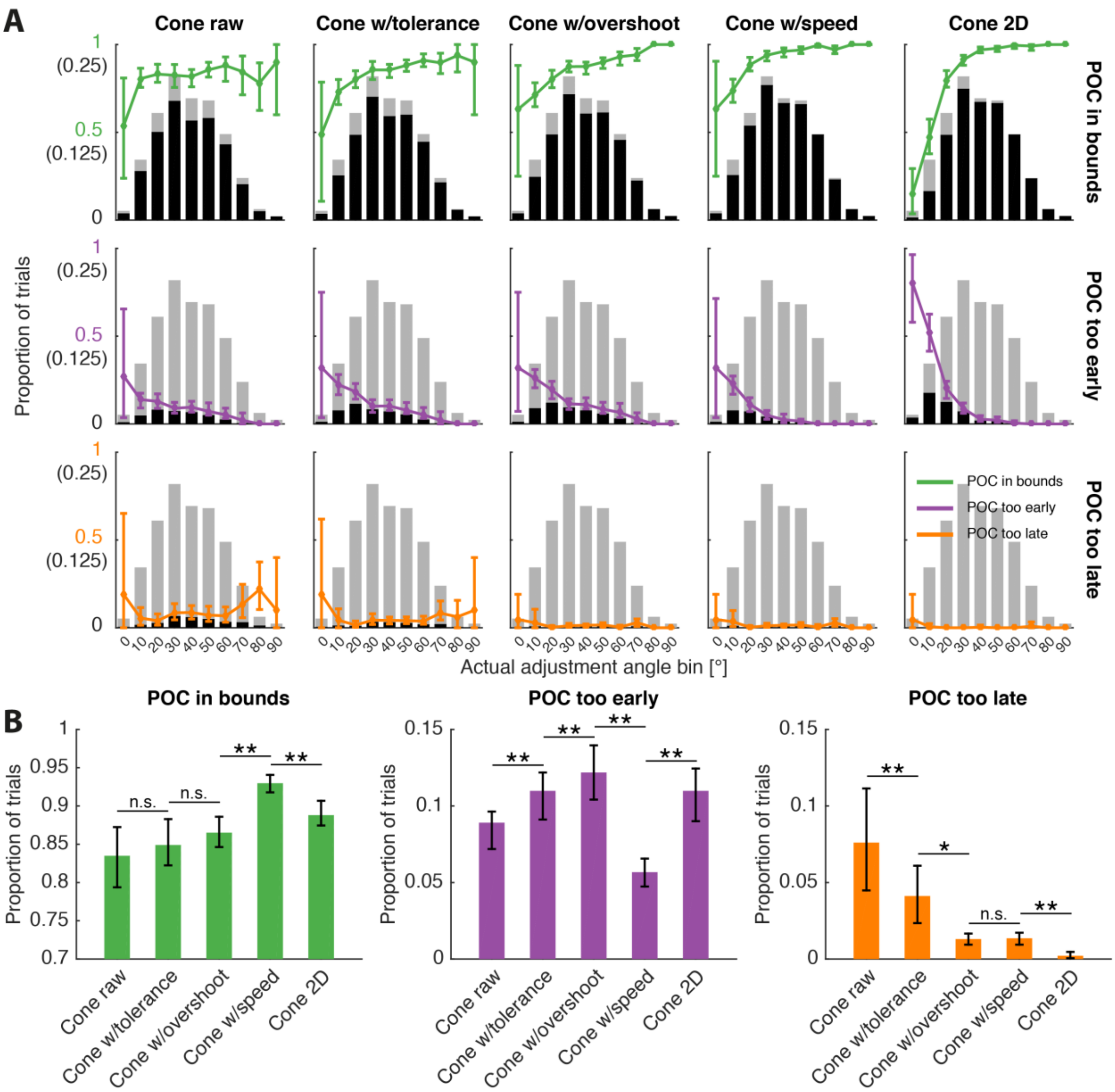


**Supplementary Figure 1-1. Influence of the different additions to the cone method on the proportion of out-of-bounds POC$^{cone}$. A:** The line graphs show the mean per-subject proportions of in-bounds (top row), too-early (middle), and too-late POC$^{cone}$ (bottom), separately for each actual adjustment angle bin. The columns (left to right) show the raw cone method, the method with added tolerance criterion, with additionally added overshoot criterion, with additionally added speed criterion (referred to as the "cone method" in the main text), and applied to the 2D projections of the trajectories. The gray bar graphs show the mean per-subject proportion of trials in each bin, out of the 240 trials per subject across all bins. Note that the gray data is repeated in each panel. The black bar graphs show the corresponding proportions of in-bounds-, too-early-, and too-late-POC$^{cone}$, respectively. The Y-axis tick marks without parentheses refer to the line graphs, the Y-axis tick marks in parentheses refer to the bar graphs. Both Y-axes start at zero. **B:** Same proportions as in A, pooled across actual adjustment angle bins. Asterisks in B are Wilcoxon signed-rank test p-values with * = $p$<0.025 ** = $p$<0.01, and n.s. = not significant. Error bars = bootstrapped (N = 2,000) 95% confidence intervals of the mean.

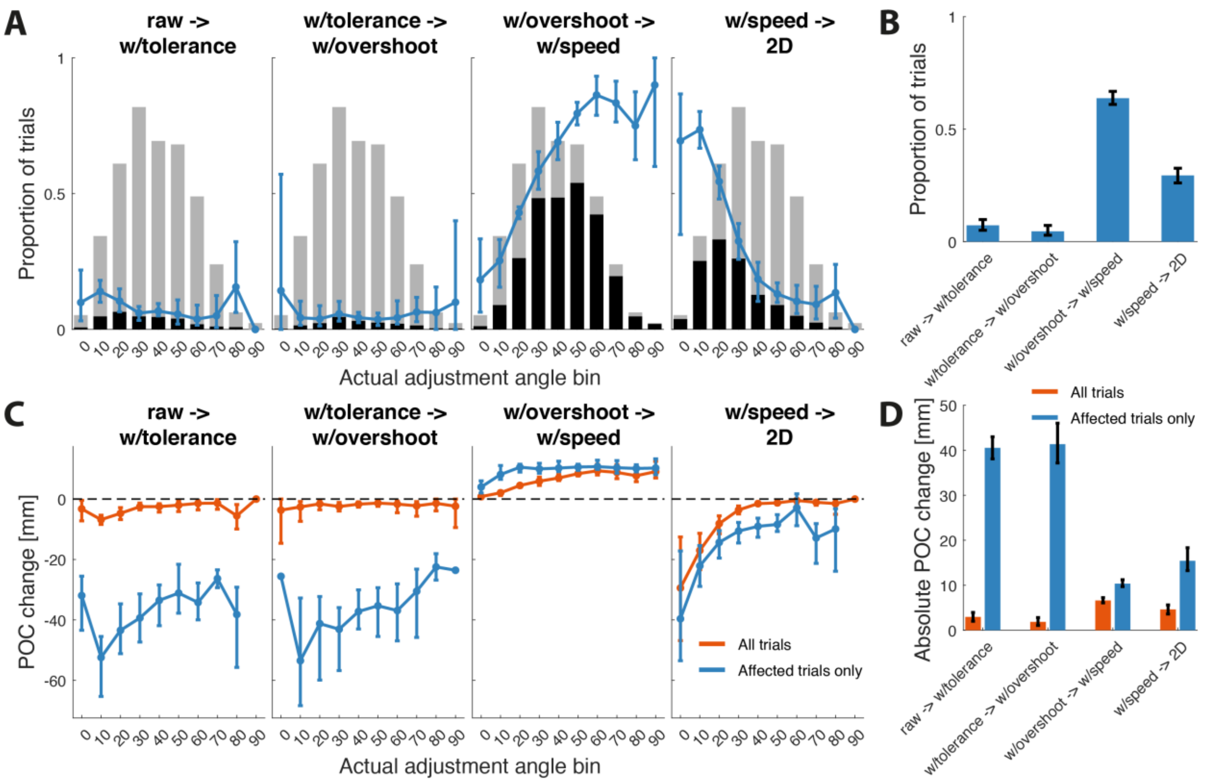


**Supplementary Figure 1-2. Influence of the different cone method additions on the POC$^{cone}$ estimates. A:** Each panel shows the mean per-subject proportion of trials affected by applying an addition to the cone method. This means, the left panel shows how many trials were affected when we applied the tolerance window to the POC$^{cone}$ obtained from the raw cone method, the mid-left panel shows how many trials were affected when we applied the overshoot adjustment to the data obtained from the cone method with applied tolerance window, and so forth. Line and bar graph conventions as in Supplementary Figure 3A. **B:** Same proportions as in A, pooled across actual adjustment angle bin. **C:** Each panel shows how the mean per-subject POC$^{cone}$ positions change due to applying an addition to the cone method (across-panel conventions as in A), either including all trials or only the affected trials. **D:** Same proportions as in C, but unsigned and pooled across actual adjustment angle bin. Error bars = bootstrapped (N = 2,000) 95% confidence intervals of the mean.

### Supplementary Information 2:

### Detailed description of the cluster-based permutation test

Cluster-based permutation tests (henceforth CP test; Maris & Oostenveld, 2007) provide a means to apply a statistical test to every sampling point of a set of continuous data while controlling for the multiple comparisons problem. Here we used the implementation by Dann, Michaels, Schaffelhofer, & Scherberger (2016) to identify from which point onward grouped trajectories towards opposite targets began to significantly branch. We interpreted this significance onset as alternative, trial-averaged measure of the point of overt commitment.

In both experiments, we used CP tests to test the lateral deviation of each set of trajectories (2D projections) against the lateral deviation of trajectories towards the opposite direction. In Experiment 1, we pooled the trajectories towards the 45° and 135° targets and 225° and 315° targets, respectively (i.e. we tested upward vs. downward trajectories). The CP test was separately applied to each grouped-pair of trajectories (i.e. Experiment 1: subject * nominal adjustment angle; Experiment 2: subject * SOA * gain versus loss trial) as follows: At each point along the trajectories, we performed a two sample t-test against the group of trajectories aimed at the opposite target. The t-tests were one-sided as we were only interested in the shift of lateral deviation towards the final target's direction. Clusters were defined as adjacent points with lateral deviation t-values that were both, significant ($p$ < 0.05) and showed an effect in the right direction (originally, when performing two-sided tests, separate clusters would be defined for positive and negative t-values). In each cluster, the t-values were summed up to create a single test statistic for each cluster ("t-sum"). We then repeated this process 1,000 times, but randomly permutated for each point whether the lateral deviation value would be labelled as belonging to movements towards their actual direction or the opposite direction. We retained the highest t-sum from each of the 1,000 iterations to create a distribution of t-sum. We then calculated the p-value for each of our original, unpermutated clusters as the ratio of t-sum distribution values that was higher than the t-sum of each respective original cluster. If this p-value was below 0.05, all time points within this cluster were considered significantly different from zero.

### Supplementary Information 3: Estimation of the actual adjustment angle (Experiment 1)

In Experiment 1, we varied the distance between starting- and via-sphere, and the lateral offset angle of the target to probe the performance of the cone method over a broad range of adjustment angles. Due to the size of the starting-, via-, and target spheres, as well as the 30° curvature tolerance within the movement corridors, the actual angle at which subjects adjusted their initial movement towards the target deviated from the task-determined nominal adjustment angles. We estimated the actual adjustment angle by fitting a tangent line each to the movement segment prior to the adjustment-to-target and after the adjustment-to-target and defined the actual adjustment angle as measured between these tangent lines (Supplementary Figure 3-1A). We determined the position of the tangent lines, using the 2D projections of the trajectories, as follows. We rotated and mirrored each trajectory such that its start- and endpoint were aligned with the X-axis and the center of the rotated via-sphere position had a positive Y-axis coordinate (Supplementary Figure 3-1B). This resulted in a positive slope for the movement portion towards the via-sphere, i.e. prior to the adjustment, and a negative slope for the movement portion towards the target, i.e. after the adjustment. We differentiated the Y-coordinates of the rotated trajectory with respect to its X-coordinates and determined the fitting points of the tangent lines as the last local maximum in the derivative prior to the sign change from positive to negative and the first local minimum after the sign change, respectively (Supplementary Figure 3-1C). This allowed us to capture the immediate movement direction tendencies prior to and after the target-oriented direction adjustments, respectively. For example, regarding the trajectory shown in Supplementary Figure 3-1 A-C, the subject moved away from the target shortly before it adjusted the movement towards the target, which led to a larger adjustment angle than would have been the case if the subject had moved in a straight line. By fitting the pre-adjustment tangent as described above, we were able to capture this deflection, thereby arriving at a more accurate actual adjustment angle estimate.

Supplementary Figure 3-1D shows that the median actual adjustment angle estimates consistently lied slightly above the nominal adjustment angles. At small nominal adjustment angles, the actual adjustment angles were larger for upward in comparison to downward targets. This is in line with the observation described in Figure 3-1C, showing that subjects often moved towards the via-sphere in an upward ark.

At small nominal adjustment angles, the spatial bias led to a larger proportion of too-early-$POC^{cone}$ for downward compared to upward targets (Supplementary Figure 3-2A). This erroneously suggests an implausibly poorer performance of the cone method for downward targets. By sorting the too-early-POC data by the actual adjustment angle instead of the nominal adjustment angle, one can visually appreciate that this effect was largely attributable to the lower actual adjustment angles towards downward targets within low nominal adjustment angle groups (Supplementary Figure 3-2B). In other words, when considering actual instead of nominal adjustment angles, it becomes apparent that the smaller the adjustment angle, the higher the risk of a too-early-POC assignment, independent of target direction.  Thus, we opted to sort the data according to actual adjustment angle instead of nominal adjustment angle, which allowed us to pool data across target directions for all subsequent analyses.

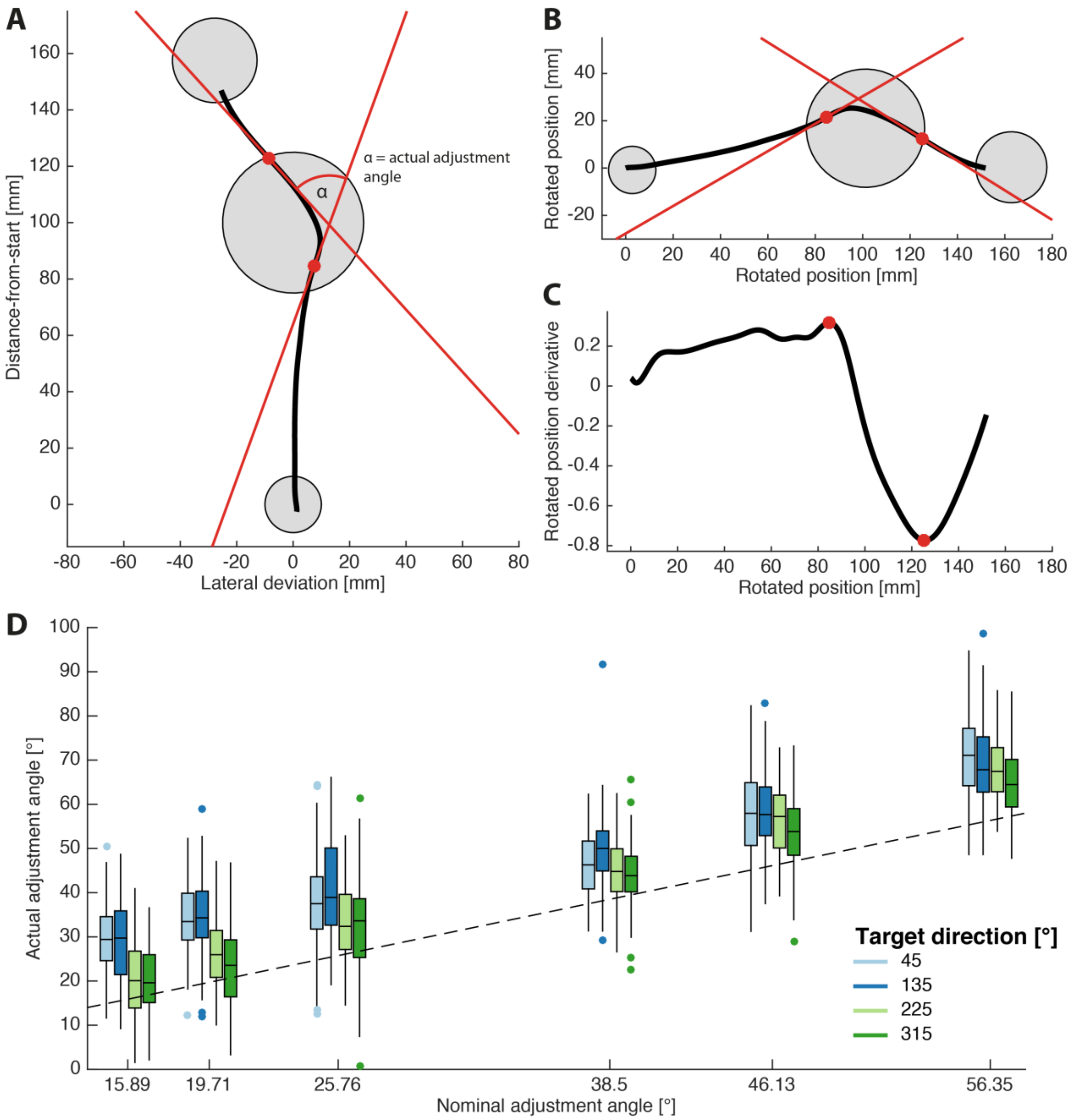


**Supplementary Figure 3-1. Estimation of the actual adjustment angle. A.** Example trajectory. The actual adjustment angle is measured between the red tangent lines which were fitted to estimate the movement direction before and after the adjustment towards the target sphere. **B.** Same trajectory as in A but rotated to align the lateral deviation coordinate of the first and last datapoint with the X-axis. If required, the rotated trajectory was additionally mirrored relative to the X-axis to ensure a positive slope before and a negative slope after the movement adjustment. **C.** Derivative of B's Y coordinates. The pre-adjustment tangent line was fitted to the trajectory point that corresponded to the derivative's local maximum closest to the center of the via-sphere. The post-adjustment tangent line was fitted to the derivative's local minimum closest to the center of the via-sphere. **D.** Resulting actual adjustment angle estimates as function of the nominal adjustment angle. Boxplots (median ± $25^{th}/75^{th}$ percentile & 1.5x interquartile range) show the aggregated single trial data across all subjects.

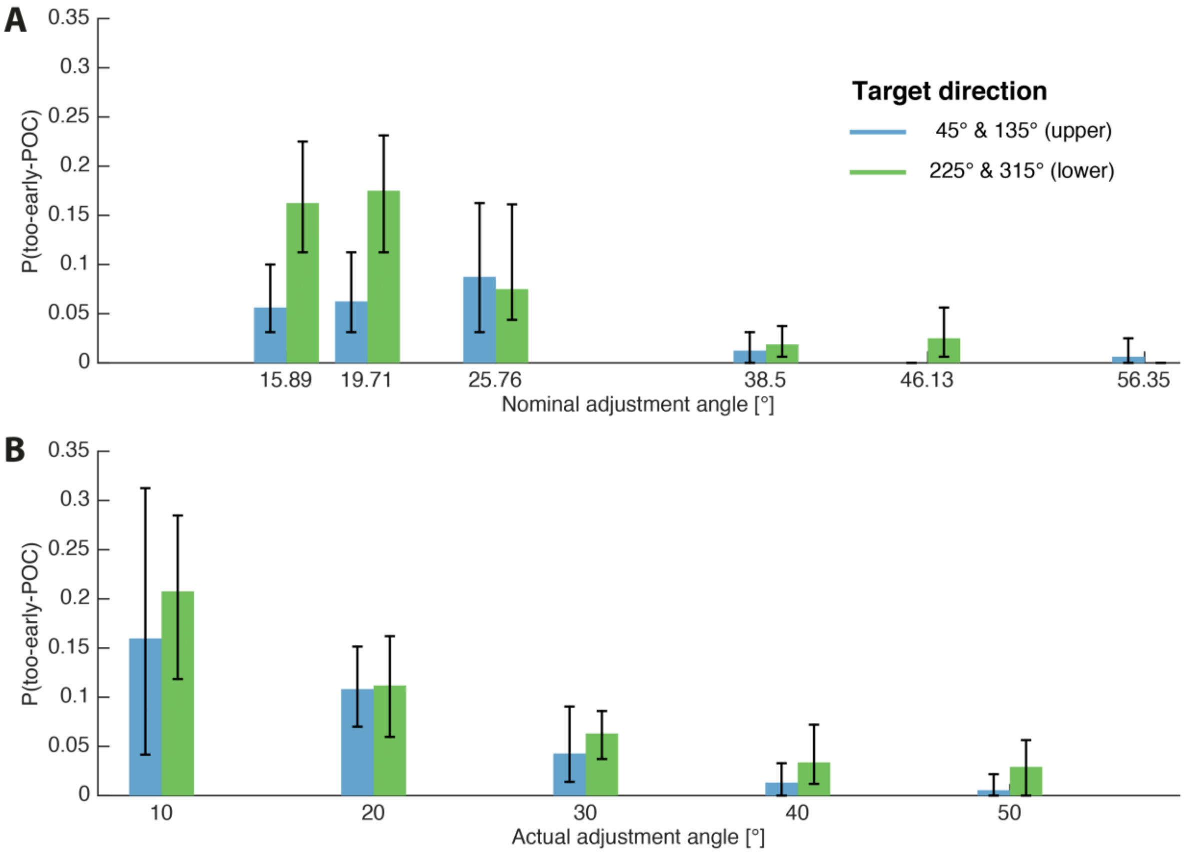


**Supplementary Figure 3-2. Influence of grouping the proportions of too-early-POC$^{cone}$ according to nominal versus actual adjustment angles. A:** Mean per-subject proportions of too-early-POC$^{cone}$ grouped according to the nominal adjustment angle. **B.** Mean per-subject proportions of too-early-POC$^{cone}$ grouped according to the actual adjustment angle. Bin width = 10°. We omitted the <10° bin (fewer than 5 trials for 7 out of 8 subjects) and all bins >50° (no too-early-POCs). Error bars = bootstrapped (N = 2,000) 95% confidence intervals of the mean.

### Supplementary Information 4: Generalized linear mixed effects model results

**Supplementary table 4-1.** Results of GLME1-1

| Dependent variable | Effect | Estimate | 95% CI | | *p* |
|---|---|---|---|---|---|
| | | | LB | UB | |
| $POC^{cone}$ in bounds | Intercept | 0.681 | 0.216 | 1.146 | .005 |
| | Actual adjustment angle | 0.061 | 0.047 | 0.075 | <.001 |
| $POC^{cone}$ too early | Intercept | -0.376 | -0.841 | 0.09 | .11 |
| | Actual adjustment angle | -0.084 | -0.102 | -0.066 | <.001 |
| $POC^{cone}$ too late | Intercept | -4.461 | -5.672 | -3.251 | <.001 |
| | Actual adjustment angle | 0.004 | -0.024 | 0.0315 | .79 |

*Note. CI* = confidence interval; *LB* = lower boundary; *UB* = upper boundary. Separate models were computed for each dependent variable.

**Supplementary table 4-2.** Results of GLME1-2

| Dependent variable | Effect | Estimate | 95% CI | | *p* |
|---|---|---|---|---|---|
| | | | LB | UB | |
| $POC^{cone}$ relative to via-sphere entry | Intercept | 10.629 | 8.307 | 12.591 | <.001 |
| | Actual adjustment angle | 0.228 | 0.187 | 0.269 | <.001 |

*Note. CI* = confidence interval; *LB* = lower boundary; *UB* = upper boundary.

**Supplementary table 4-3.** Results of GLME1-3

| Dependent variable | Effect | Estimate | 95% CI | | *p* |
|---|---|---|---|---|---|
| | | | LB | UB | |
| POC$^{cone}$ relative to via-sphere entry | Intercept | 14.444 | 12.153 | 16.736 | <.001 |
| | Nominal adjustment angle | 0.16 | 0.104 | 0.215 | <.001 |
| POC$^{CP}$ relative to via-sphere entry | Intercept | 30.367 | 23.52 | 37.215 | <.001 |
| | Nominal adjustment angle | -0.217 | -0.383 | -0.05 | .012 |

*Note. CI* = confidence interval; *LB* = lower boundary; *UB* = upper boundary. Separate models were computed for each dependent variable.

**Supplementary table 4-4.** Results of GLME2-1

| Dependent variable | Effect | Estimate | 95% CI | | *p* |
|---|---|---|---|---|---|
| | | | LB | UB | |
| TOC$^{cone}$ at movement start | Intercept | 0.121 | -0.584 | 0.825 | .73 |
| | Value cue SOA | -0.006 | -0.009 | -0.002 | .002 |
| TOC$^{cone}$ before value cue | Intercept | -7.867 | -9.23 | -6.504 | <.001 |
| | Value cue SOA | 0.0161 | 0.012 | 0.02 | <.001 |
| All early TOC$^{cone}$ | Intercept | 0.012 | -0.675 | 0.698 | .97 |
| | Value cue SOA | -0.005 | -0.008 | -0.001 | .005 |

*Note. CI* = confidence interval; *LB* = lower boundary; *UB* = upper boundary. Separate models were computed for each dependent variable.

**Supplementary table 4-5.** Results of GLME2-2

| Dependent variable | Effect | Estimate | 95% CI | | *p* |
|---|---|---|---|---|---|
| | | | LB | UB | |
| $TOC^{cone}$ | Intercept | 436.1 | 387.4 | 484.8 | <.001 |
| | Value cue SOA | 0.525 | 0.4041 | 0.646 | <.001 |
| | $Frame_{gain/loss}$ | -55.215 | -82.393 | -28.038 | <.001 |
| | SOA × frame | 0.179 | 0.064 | 0.293 | 0.003 |
| $TOC^{CP}$ | Intercept | 436.42 | 375.51 | 497.33 | <.001 |
| | Value cue SOA | 0.608 | 0.426 | 0.789 | <.001 |
| | $Frame_{gain/loss}$ | -64.375 | -102.41 | -26.343 | .001 |

*Note. CI* = confidence interval; *LB* = lower boundary; *UB* = upper boundary. Separate models were computed for each dependent variable. Main effects were obtained from models without interaction effect.
