## Supplementary figures and images for "The Cone Method: Inferring Decision Times from Single-Trial 3D Movement Trajectories in Choice Behavior"

### Supplementary Figure 1-1

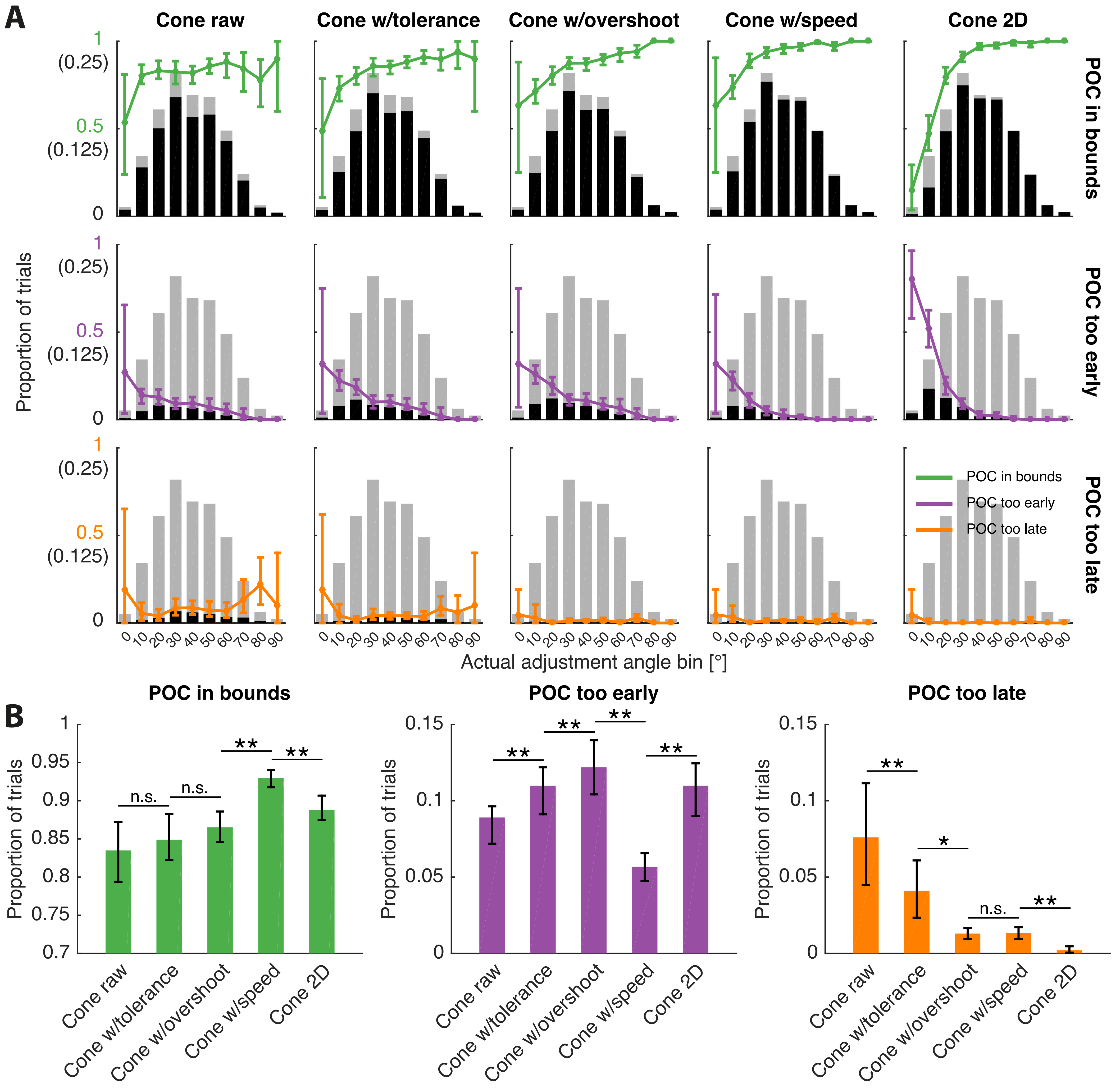

### Supplementary Figure 1-2

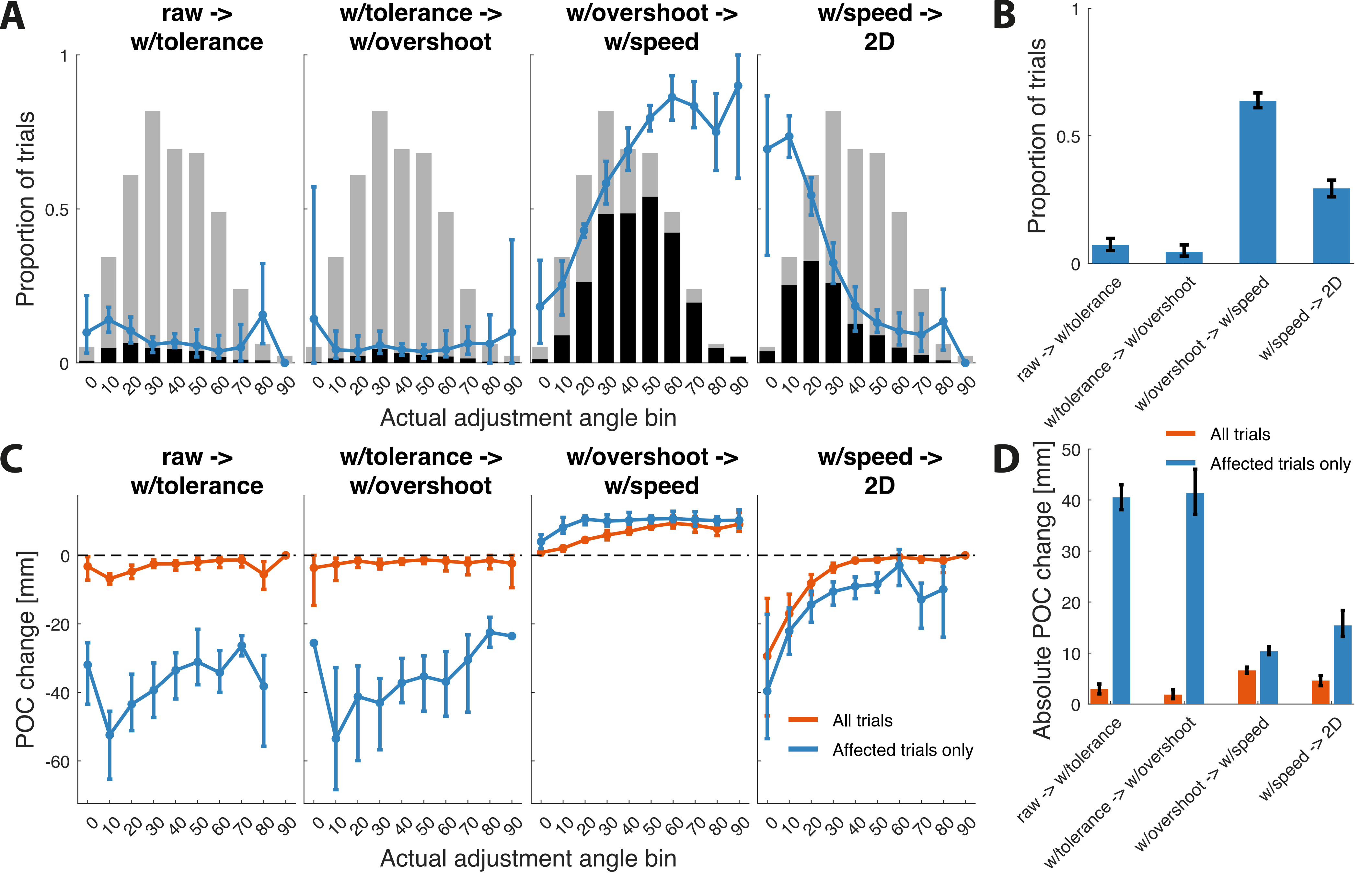

### Supplementary Figure 3-1

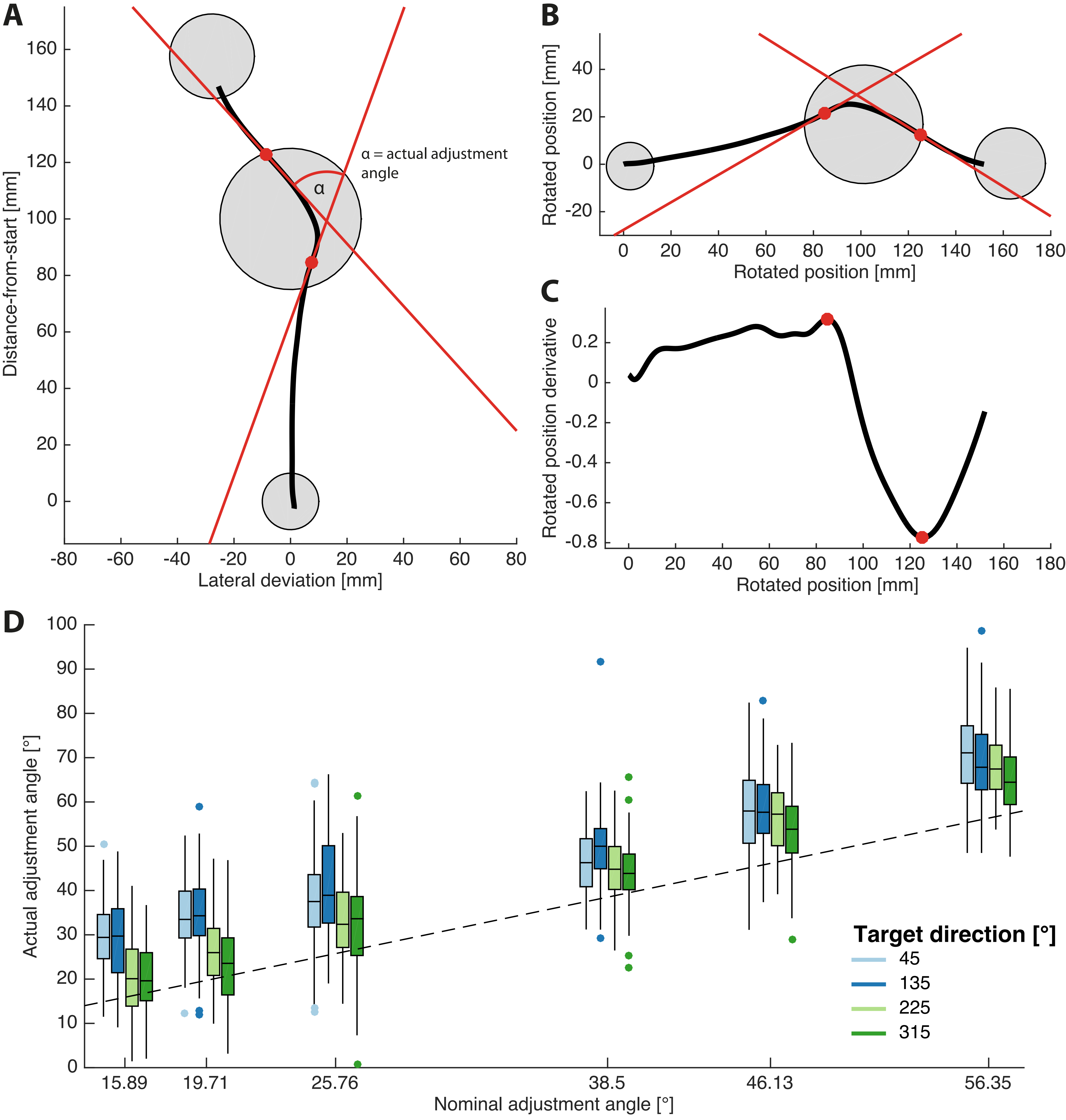

### Supplementary Figure 3-2

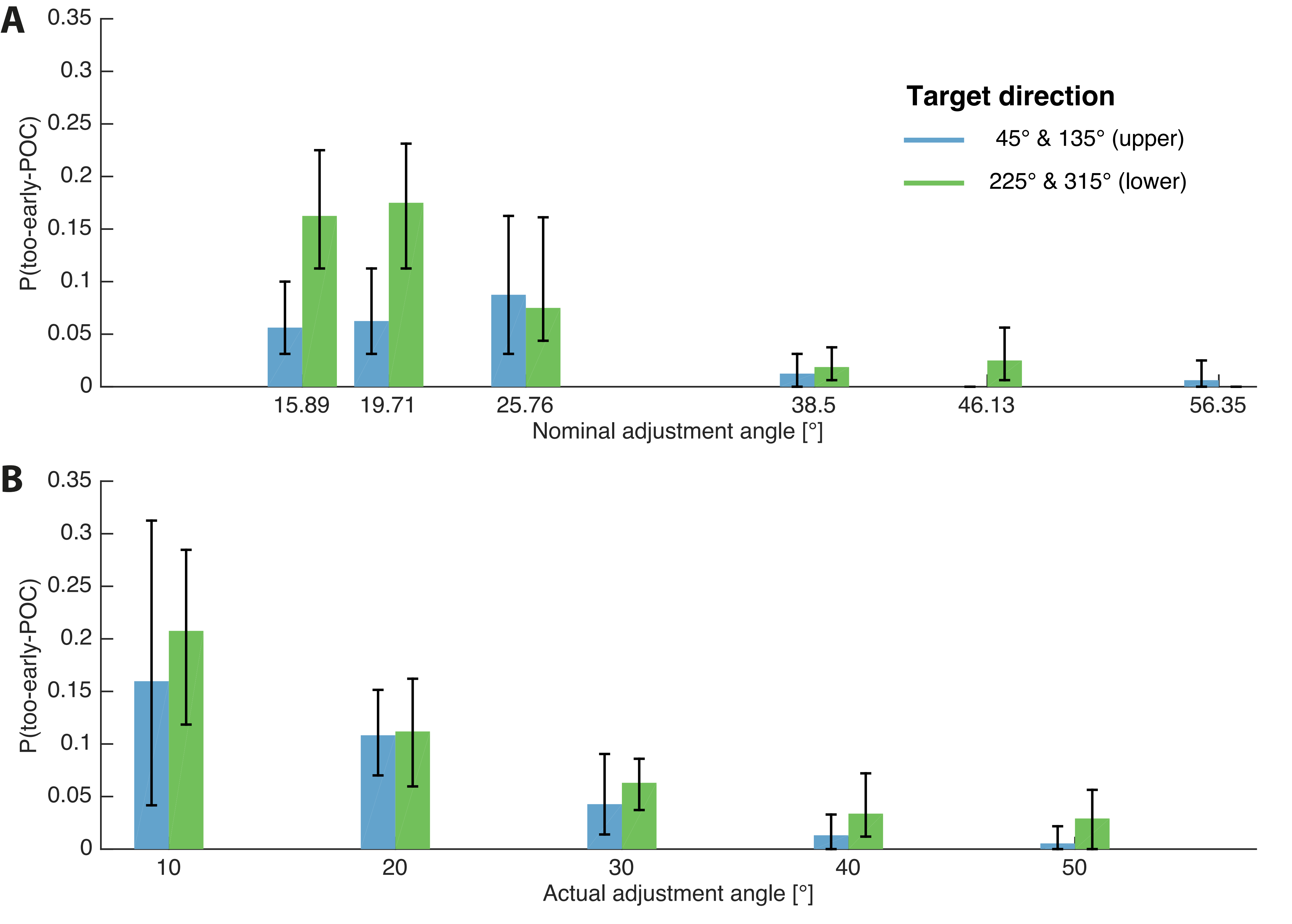
